# Can interpretability and accuracy coexist in cancer survival analysis?

**DOI:** 10.1101/2025.04.11.648380

**Authors:** Piyush Borole, Tongjie Wang, Antonio Vergari, Ajitha Rajan

## Abstract

Survival analysis refers to statistical procedures used to analyze data that focuses on the time until an event occurs, such as death in cancer patients. Traditionally, the linear Cox Proportional Hazards (CPH) model is widely used due to its inherent interpretability. CPH model help identify key disease-associated factors (through feature weights), providing insights into patient risk of death. However, their reliance on linear assumptions limits their ability to capture the complex, non-linear relationships present in real-world data. To overcome this, more advanced models, such as neural networks, have been introduced, offering significantly improved predictive accuracy. However, these gains come at the expense of interpretability, which is essential for clinical trust and practical application. To address the trade-off between predictive accuracy and interpretability in survival analysis, we propose ConSurv, a concept bottleneck model that maintains state-of-the-art performance while providing transparent and interpretable insights. Using gene expression and clinical data from breast cancer patients, ConSurv captures complex feature interactions and predicts patient risk. By offering clear, biologically meaningful explanations for each prediction, ConSurv attempts to build trust among clinicians and researchers in using the model for informed decision-making.

## 1 Introduction

Survival analysis is essential for estimating the time until events such as death, relapse, or recovery occur. It lays the groundwork for assessing disease severity and understanding the factors that influence patient outcomes [1, 2]. Traditional methods like the Cox Proportional Hazards (CPH) model, a linear approach, have been widely used due to their straightforward interpretability and robust ability to estimate hazard ratios (a type of relative risk) [2, 3]. However, with the advent of high-dimensional data such as RNA sequencing (RNA-seq), these linear models face significant limitation in achieving high performance.

RNA expression data measure gene activity levels and play a critical role in survival research by revealing dysregulated genes associated with disease progression. The complex and non-linear relationships inherent in high-dimensional RNA-seq data challenge the CPH model’s ability to accurately extract meaningful features. This limitation results in less precise predictions compared to more advanced machine learning models that can capture these intricate patterns. However, CPH, being a linear model, possesses inherent interpretability, allowing the importance of each input feature to be inferred from its feature weights. As illustrated in Figure 1, such a linear model offers high interpretability (on the x-axis) but low accuracy (on the y-axis).

**Fig. 1.**
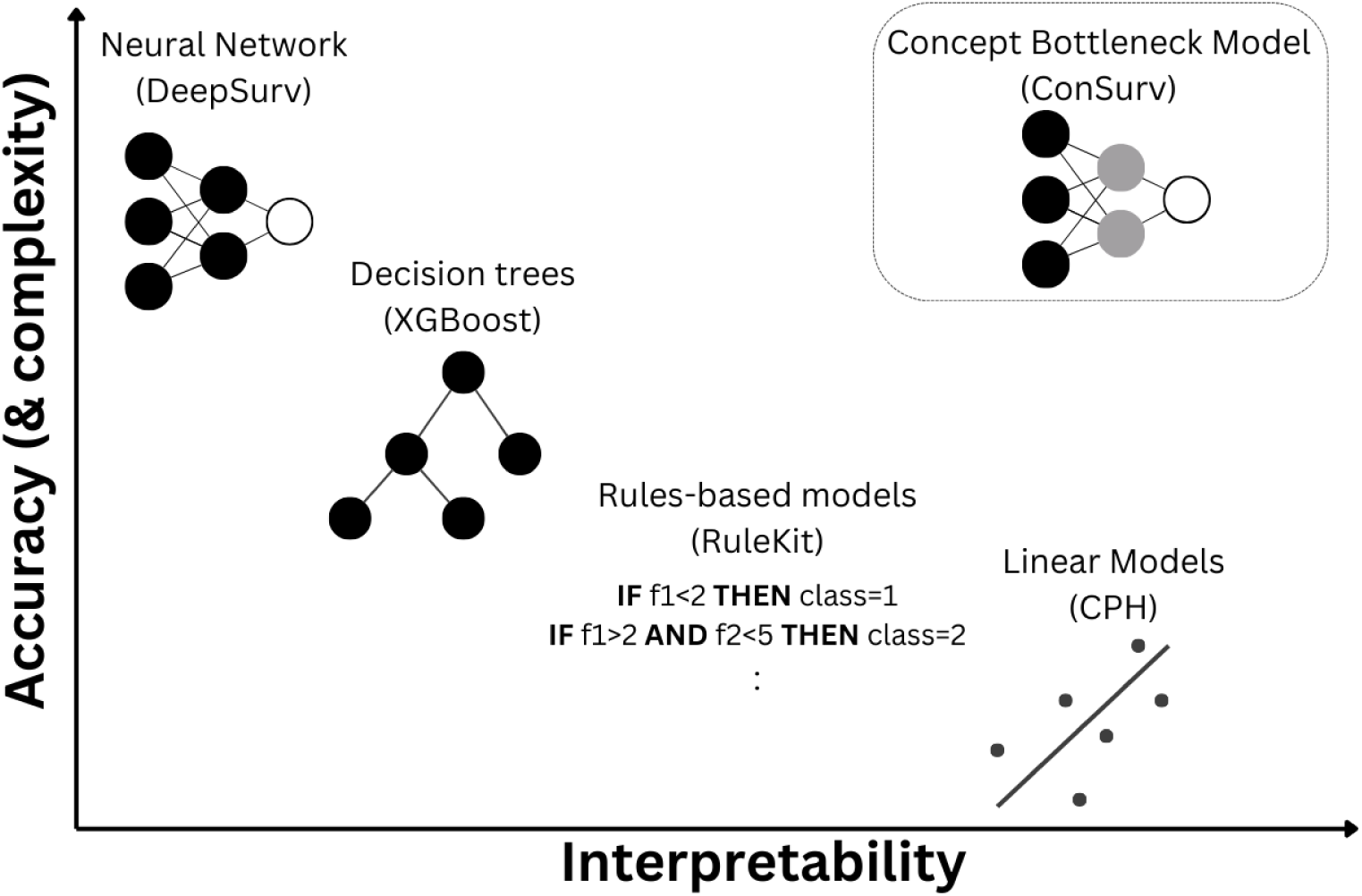
Accuracy and Interpretability trade-off. Linear and rule-based models are inherently interpretable due to their explicit rules and feature weights used for prediction. However, they often exhibit lower performance compared to more complex, black-box models such as neural networks. As complexity of models increase, they tend to improve in accuracy, but this often comes at the cost of interpretability, as their decision-making processes become more difficult to understand. Tree-based algorithms, while offering transparent decision pathways, can also become challenging to interpret due to their large number of decision nodes and depth of the tree. A concept bottleneck model, on the other hand, is a gray-box model that strikes a balance between accuracy and interpretability. It consists of two parts: a black-box component that predicts human-interpretable “concepts” and a second part that uses these concepts exclusively to make the final prediction, typically through a simple (often linear) model.

At the other extreme, emerging non-linear neural network models such as DeepSurv [4], offer improved accuracy by capturing the complexity of the data but provide very low interpretability. This lack of interpretability makes it challenging for clinicians and researchers to derive meaningful causal relationships between input features and risk predictions. This is illustrated in Figure 1, where the neural network ranks high on accuracy dimension but very low on interpretability dimension. The opacity of these models and the technical expertise required for their interpretation lead to a trust gap, preventing their adoption in clinical settings. Current research on interpretability of these neural networks for survival analysis focuses on post-hoc explanations which involve applying additional models to ascertain contribution of each feature (called explanations) to the predictions of these black-box models. For example, SurvLIME [5] uses the Local Interpretable Model-agnostic Explanations (LIME, [6]) framework, while SurvSHAP [7] and AUTOSurv [8] apply SHapley Additive exPlanations (SHAP, [9]). Both LIME and SHAP are perturbation-based models that require multiple evaluation passes over the model to explain a single prediction. Furthermore, empirical evidence demonstrates that these models could fail to assign importance to relevant features and produce explanations that are unfaithful [10]. These limitations make perturbation-based models not only computationally expensive but also, more importantly, unreliable [11, 12]. Other approaches, utilize DeepLIFT [13, 14] and backpropagation [15] to quantify the contribution of input features to the risk prediction. However, these gradient-based methods are highly sensitive to noise and non-linearities, leading to explanations that are often unreliable and unfaithful to the models they aim to interpret [16]. Relying on such explanations can lead to incorrect conclusions about the data, ultimately resulting in a trust deficit. The Cox-nnet model [17], a single-layer perceptron, takes an interesting approach to interpretability by capturing biologically relevant functions (such as the p53 pathway) in its hidden nodes, which serve as surrogate features for predicting patient survival. However, their approach of quantifying the contribution of each input to every node of the network quickly becomes infeasible as deeper networks are required to improve accuracy.

Tree-based models, such as XGBoost [18, 19], provide a partial solution by better balancing transparency and accuracy than deep learning models as seen in Figure 1. They offer transparency by using features as nodes within the trees, which could be individually examined, while also capturing non-linear relationships more effectively than linear models such as CPH. However, when applied to high-dimensional data such as RNA-seq, the trained trees can have vast number of decision nodes making them overwhelmingly complex. This growth in complexity inherently hinders interpretability.

As seen in the above examples of linear, tree-based, and neural network models, there is a trade-off between performance and interpretability, which is illustrated as accuracy-interpretability trade-off in Figure 1. This trade-off suggests that as model complexity increases, accuracy improves, but interpretability declines. An ideal model should balance both aspects to ensure usability and trustworthiness.

To balance the accuracy-interpretability trade-off in survival prediction, we propose a gray-box approach using concept bottleneck models [20]. These models consist of two parts: one predicts human-interpretable concepts, and the other—typically a linear model—uses these concepts for final predictions. We apply this approach to risk prediction on RNA-seq and clinical data for breast cancer and demonstrate performance comparable to the existing models. We also show that the key concepts can differentiate high- and low-risk patients demonstrating the usefulness of these concepts. Finally, we assess the stability of these concepts by evaluating their importance and robustness by examining the consistency in extracted concepts across multiple runs.

Our model, ConSurv, inherently embeds interpretability as a core component by aggregating genetic and clinical features into concepts, providing humanly understandable and biologically meaningful insights.

## 2 Results

In the results section, we evaluate the models based on two key dimensions: performance in predicting LogRisk and ability in explaining model decision-making (i.e. interpretability). The existing models assessed in this study are CPH, XGBoost, and DeepSurv that are popular and widely used in survival analysis and covers the spectrum on accuracy-interpretability tradeoff of Figure 1. Performance is measured using the Concordance Index (or CI, see Section 4.4).

The result section begins with description of the dataset (Section 2.1, TCGA breast cancer data) used in this study. Sections 2.2 and 2.3 present the performance evaluation of existing models and our proposed ConSurv framework. Finally, sections 2.4 to 2.6 focus on model interpretability. We begin by outlining the interpretability aspects of the linear CPH, the black-box DeepSurv (using SHAP), and the gray-box ConSurv model. This is followed by an exploration of the extracted concepts, assessing their alignment with biological insights, as well as their stability and robustness. Table 1 provides a summary of the models used in this study and their interpretability.

**Table 1.** Survival Analysis Models And Their Interpretability.

| Model | Type | White-/<br>Black-box | Interpretability |
| --- | --- | --- | --- |
| Cox Proportional Hazards | Linear Model | White-box | Feature Weights |
| XGBoost | Non-Linear<br>(Decision Trees) | White-box | Decision Tree, Post-hoc |
| DeepSurv | Non-Linear<br>(Neural Network) | Black-box | Post-hoc |
| **ConSurv** | **Non-Linear**<br>**(Concept Bottleneck Model)** | **Gray-Box** | **Learned Concepts** |

### 2.1 Data

In this study, we utilized breast cancer RNA-seq (RNA profiling table) data and corresponding clinical information obtained from The Cancer Genome Atlas (TCGA) through the Genomic Data Commons (GDC) portal.

All data used in this study are de-identified and publicly available, eliminating the need for additional ethical approval. The study complies with the data usage policies of TCGA and GDC, which allow the use of the data for research purposes.

RNA profiling table, contains more than 60,000 gene expression values per patient. Given the limitations of physical memory and training time constraints, we opt for the variance threshold method to filter out genes that contain less information. Therefore, we rank the index of dispersion for each gene and select the top 1000 highest-ranked genes. In addition to these 1000 genes, we added 8 clinical features associated with patients to the data set. These clinical features were one hot encoded. In total, there were 1209 patients with 1056 features in our final dataset. Each patient entry was accompanied with survival time (*t*) and event indicator (*e*, which indicates if event, here death, occurred or not).

### 2.2 Non-linear models demonstrate significantly better performance than linear model

The purpose of risk prediction is to stratify patients such that those at higher risk would have lower survival time or probabilities. To predict this risk (predicted as LogRisk), we used 1000 genes from RNA-seq data and eight associated clinical features for 1209 breast cancer patients. Model performance was evaluated using the Concordance Index (CI), as described in Section 4.4. In our study, we utilized widely used models - CPH, XGBoost, and DeepSurv, from the linear, tree-based, and neural network categories, respectively, as shown in Table 1 (and Figure 1). CPH is a linear model that provides association between the survival time of patients and the features. XGBoost is tree-based modeling approach used for survival regression that predicts survival time [19]. DeepSurv is a neural network that predicts LogRisk using Cox-loss function (See Section 4.6) [4]. In addition to these, we trained a multi-layer perceptron (MLP) to establish baseline parameters (such as network depth) for our concept bottleneck model. This base MLP was subsequently transformed to develop our ConSurv model as described in Section 4.2.

Data was split ten times using different seeds into training, validation and test sets (70%, 15% and 15%). Training and validation sets were used for hyperparameter tuning (See Section 4.3). Performance (i.e. CI) of the best models was evaluated using the ten test sets. The linear CPH model exhibits the lowest median performance (median CI 0.65), while the black-box DeepSurv (median CI 0.75) achieves the highest median performance, with XGBoost (median CI 0.69) falling in between. The MLP demonstrates performance comparable to DeepSurv (median CI 0.73, p-value*>*0.05, see Supplemental Table 2). Figures 2a,b plots the CI for the above described models. For XGBoost, hyperparameter tuning revealed that the highest performance was achieved with a tree depth of 10. During each run, 330–390 trained trees were generated. XGBoost also exhibits the highest variance in performance, with the lowest CI being 0.53 and the highest CI reaching 0.87 as seen in Figure 2a. We pruned the trained XGBoost trees to depths ranging from 1 to 10 and found that pruning upto a depth of 5 resulted in a marginal loss in performance (within 5% of the model with a depth of 10), as shown in Figures 2c,d. An illustration of tree pruning is in Supplemental Figure 1 which shows a tree with original depth 10 pruned upto depth 1. Across the runs, only 14 features (of 1056) were consistently used by XGBoost, as shown in Table 2. This low overlap of common features could explain the high variability in its performance. Nevertheless, the most commonly used features (predominantly genes) are known to be associated with breast cancer survival, and their references are provided in Table 2.

**Fig. 2.**
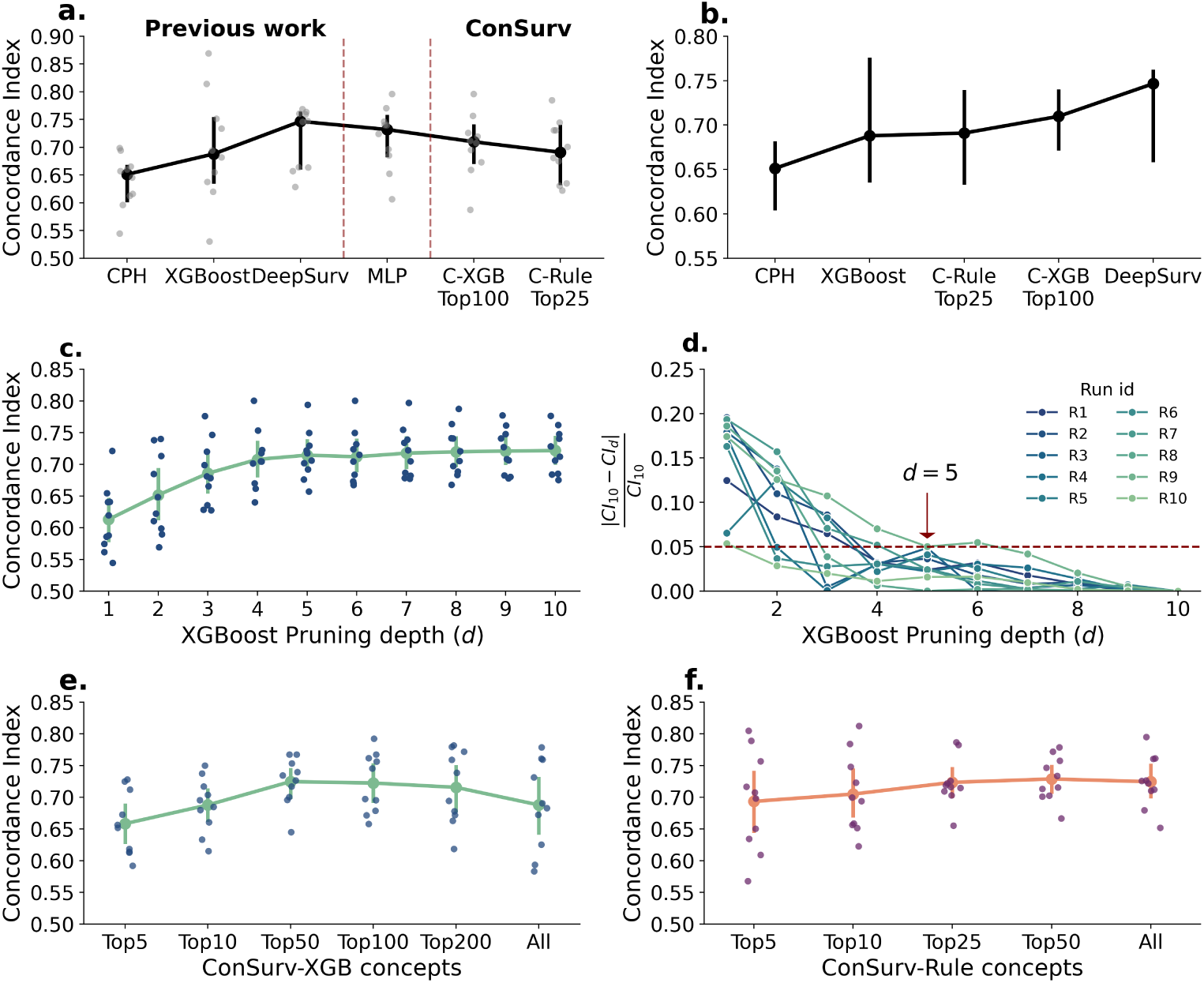
Performance of the survival models. **a.** Concordance Index or CI on test sets for Cox Proportional Hazards (CPH), XGBoost, DeepSurv, MLP and ConSurv models. Top 100 concepts were used for ConSurv-XGB (C-XGB Top100) and top 25 concepts for ConSurv-Rule (C-Rule Top25). **b.** Shows a upward CI trend as models progress from a fully interpretable to black-box. **c.** demonstrates the impact of pruning XGBoost trees to depths of 1–9 (with 10 being the original depth). **d.** Notably, the XGBoost performance with pruning depths of 5–9 remains within 5% of the original CI for depth 10 across all ten runs. **e.** Validation CI after training C-XGB using the top 5, 10, 50, 100, and 200 concepts ranked by weights in ConSurv-XGB using all concepts. **f.** Validation CI after training C-Rule model with top 10, 25, and 50 concepts ranked by weights from ConSurv-Rule using all concepts. (For all plots, errorbars: confidence interval at 95%, estimator: median)

**Table 2.**
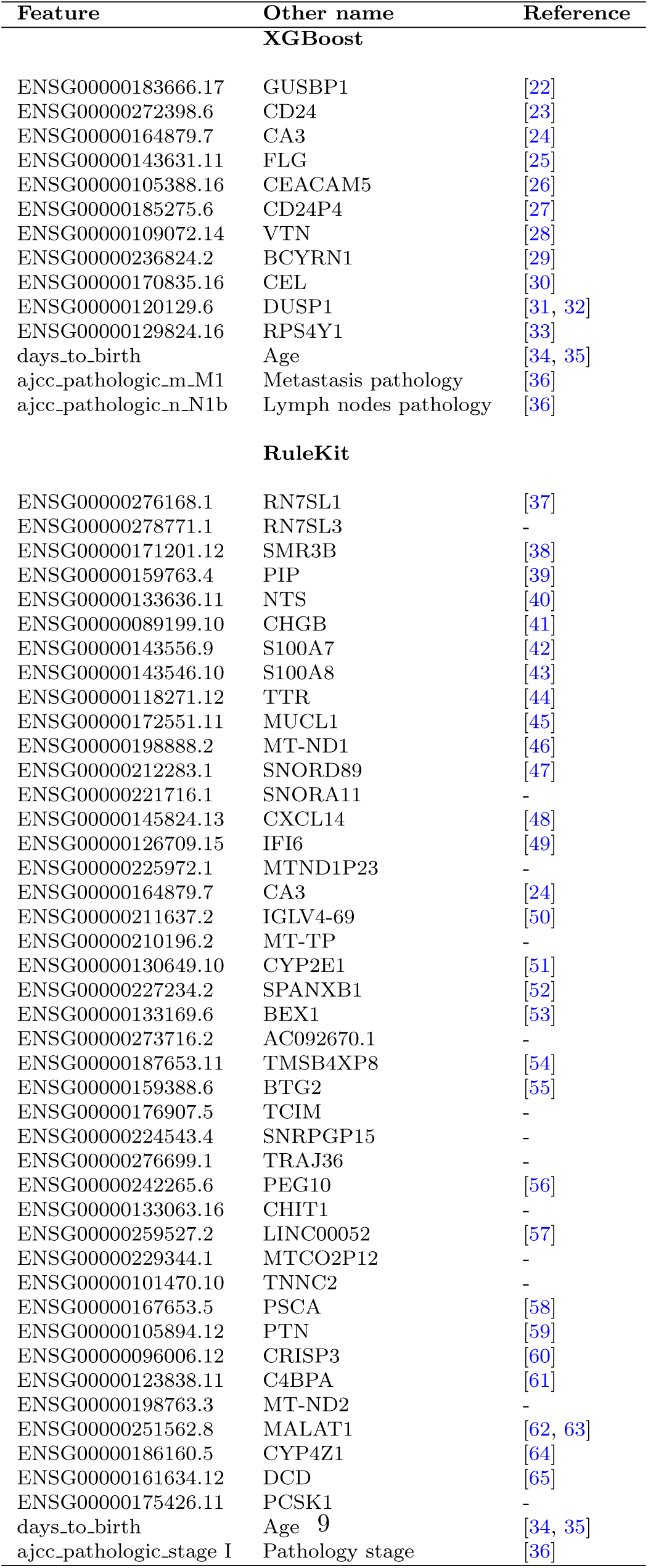
Common features across 10 different runs from XGBoost and RuleKit (Genes converted using https://www.biotools.fr/)

#### Survival classification using Rule-based model

As an additional means for pattern extraction used in our study, we trained a contrast set rule-based model (See Section 4.1), a form of association rule mining. The Contrast Set Rule-based model identifies relationships between features to cluster data points into classes using an ‘IF-THEN’ formulation. RuleKit is an implementation of this model which was used in our study. We transformed our problem into a classification task by categorizing patients into three groups based on survival times as standard practice: *<* 5 years, 5–10 years, and *>* 10 years.

The classification AUROC were 0.59 (*<* 5 years), 0.6 (5–10 years) and 0.55 (*>* 10 years) for the three classes. Each run produced between 61–83 rules, with median rule length (or number of features per rule) of 21. About 75% of rules from each run belonged to class *<* 5 survival years, 20% belonged to class 5–10 survival years and remaining 5% belonged to class *>* 10 survival years. Across runs, 44 features were consistently used by RuleKit, as shown in Table 2.

To determine whether these consistently relevant common genes of XGBoost and RuleKit play a central role in survival outcomes, we performed Gene Set Variation Analysis (GSVA) [21]. GSVA enables a non-parametric assessment of gene set enrichment. The dataset specifications for running GSVA are provided in Section 4.7. Figures 3a,b illustrate the survival stratification based on common genes for both models.

**Fig. 3.**
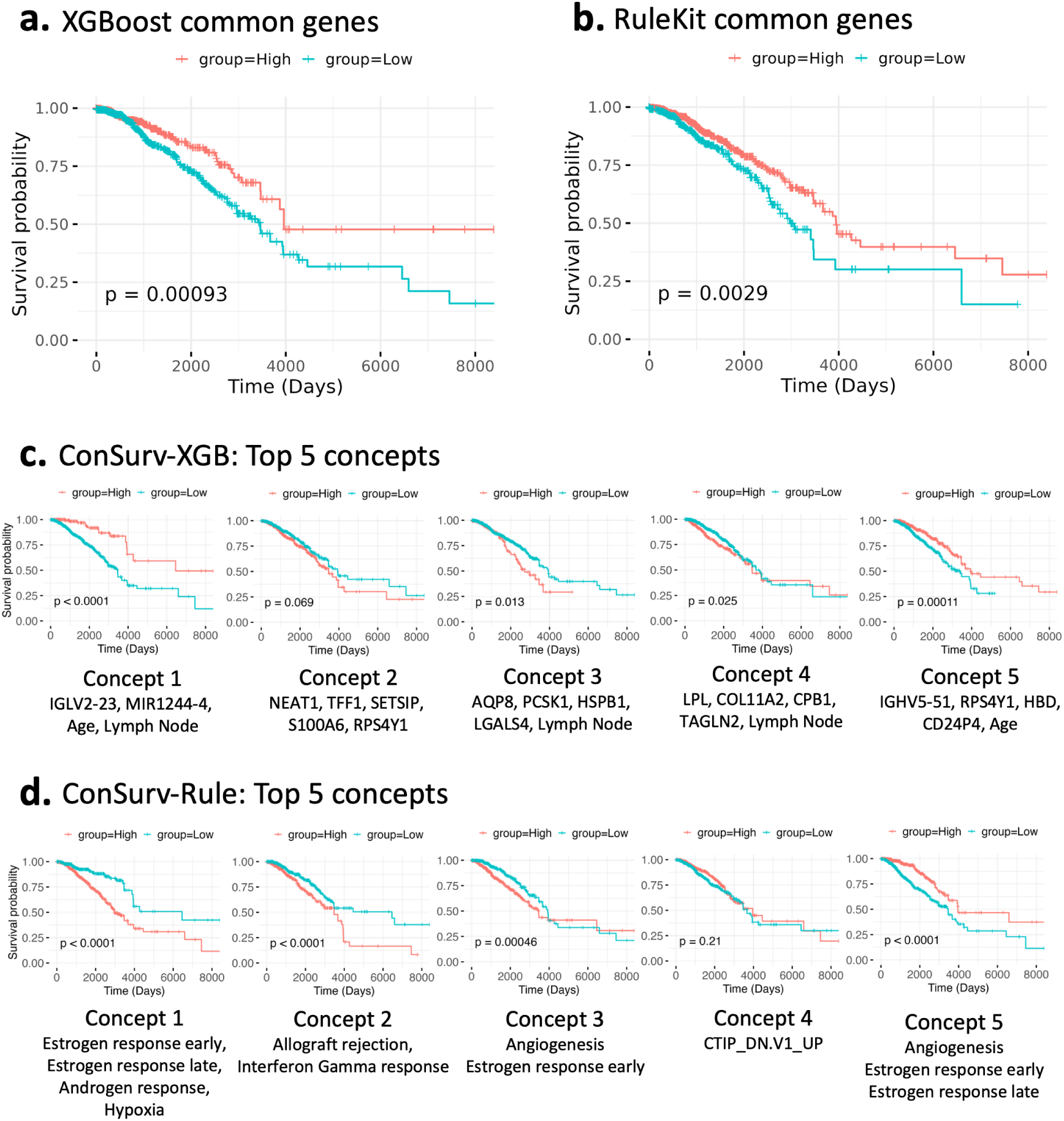
Kaplan-Meier (KM) survival plots. **a.** The KM plot for common genes selected across runs (listed in Table 2) for XGBoost shows a high level of stratification between high- and low-risk groups (*p* = 0.00093). **b.** The KM plot for common genes selected across runs (listed in Table 2) for RuleKit also demonstrates significant stratification between high- and low-risk groups (*p* = 0.0029). **c.** The KM plots for the top 5 concepts produced by the ConSurv-XGB model show significant separation between low- and high-risk groups for all but one concept (concept 2). The features forming the concept are listed under respective KM plot. **d.** Similarly, the KM plots for the top 5 concepts produced by the ConSurv-Rule model demonstrate significant separation between low- and high-risk groups for all but one concept (concept 4). The processes obtained through ORA for each concept are listed below the concept.

For XGBoost and RuleKit, a significant stratification between high-risk and low-risk groups is observed (*p* = 0.00093 and *p* = 0.0029, respectively). GSVA further reveals that the common genes identified by XGBoost are associated with the Gene Ontology (GO) molecular function GO:0005198 (i.e., structural molecule activity), which is relatively nonspecific. In contrast, the common genes identified by RuleKit are linked to a more specific biological process—programmed cell death (GO biological process GO:0012501). The ability of both models to leverage common genes for survival stratification underscores their effectiveness in identifying core biological features that contribute to predictive performance.

### 2.3 ConSurv enhances interpretability while maintaining competitive performance

Our proposed model ConSurv, is a concept bottleneck model that predicts risk as described in Equation (5) with human interpretable concepts as an intermediate prediction. The architecture is based on the bilayered MLP model with the second layer modified to capture predetermined concepts as illustrated in Figure 4a. In our framework, concepts are defined as sets of grouped features extracted either from trained trees generated by XGBoost or from rules learned using the Contrast Set Rule-based model (RuleKit), as described in Section 4.2. The two ConSurv models are named as ConSurv-XGB-all and ConSurv-Rule-all respectively. The ‘-all’ indicates use of all concepts extracted by XGBoost or RuleKit.

**Fig. 4.**
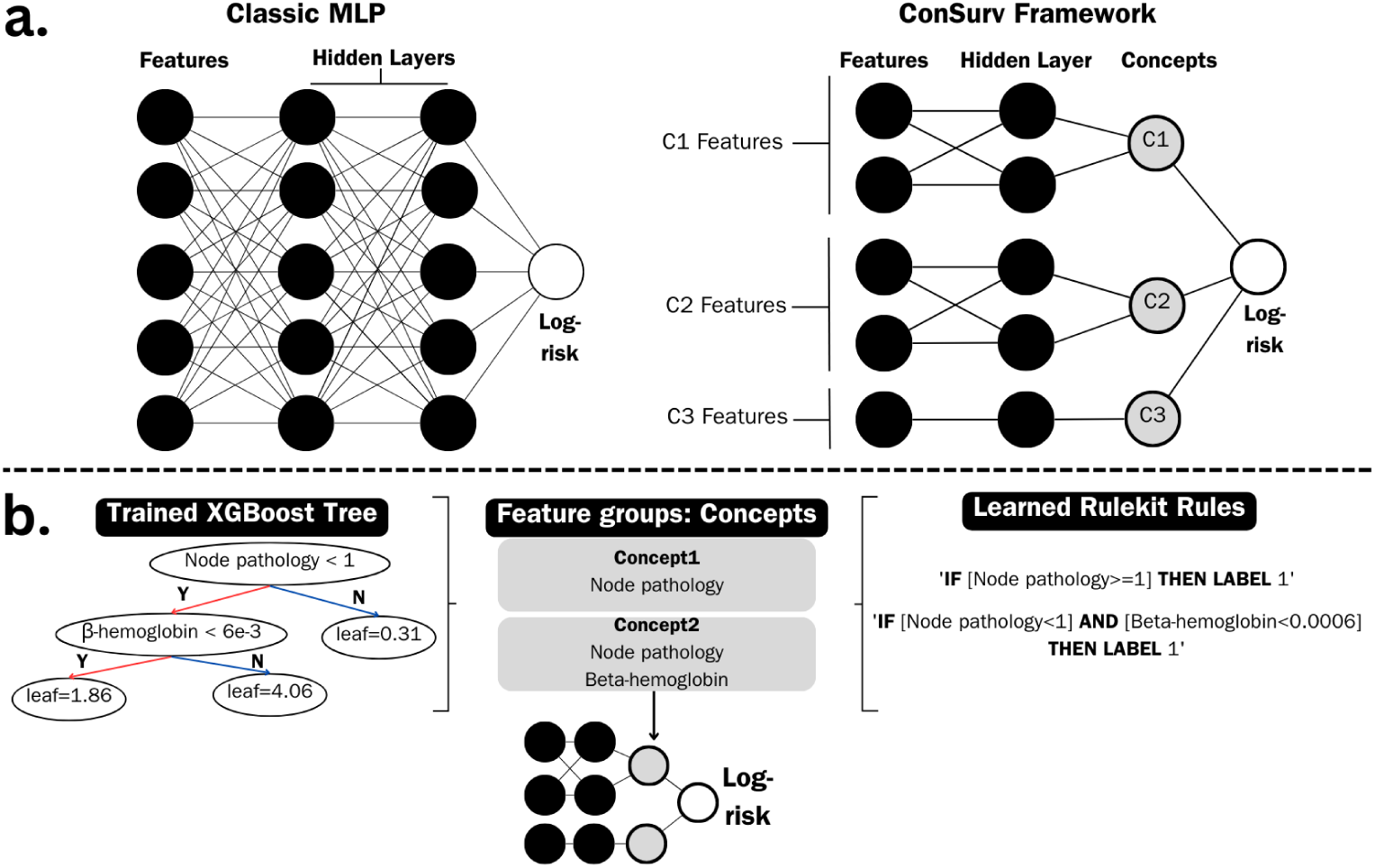
ConSurv illustration. **a.** The network on the left depicts a typical multi-layered perceptron model with two hidden layers. This densely connected architecture provides no explanation for its decision-making process. The network on the right illustrates our concept bottleneck model, ConSurv. ConSurv consists of one hidden layer followed by a concept layer, where each concept receives input only from the features associated with that concept. The final LogRisk prediction is computed as a linear combination of these concepts. **b.** Concepts are extracted using either a trained XGBoost model or rules from the RuleKit model. For RuleKit, each rule’s unique features are grouped as a concept. For XGBoost, concepts are extracted at each tree depth.

Each run of XGBoost (pruned to depth 5) generated between 540 and 770 concepts. However, having such a large number of concepts can hinder the model’s interpretability. To address this, we investigated reducing the number of concepts while maintaining performance. We first ranked the concepts from the trained ConSurv-XGB model based on their absolute weights. Then, we trained the model using only the top 5, 10, 50, 100, and 200 concepts. Supplemental Figure 2 illustrates the ranked concepts in descending order of absolute weight across different runs. As shown in Figure 2e, the validation set CI improves with increasing number of concepts upto 100 but plateaus (and even slightly declines) beyond that, when using all concepts. Based on this, we consider ConSurv-XGB with the top 100 concepts (hereafter referred to simply as ConSurv-XGB or C-XGB Top100) to be the best model in our analysis, achieving a median test set CI of 0.71. This approach reduces the number of concepts by approximately 80–88%, significantly simplifying the model while maintaining performance.

Each run of RuleKit generated between 61 and 83 concepts, with an average of 25–30 features per concept. Similar to ConSurv-XGB, we evaluated models using the top 5, 10, 25, 50, and all available concepts. Supplemental Figure 3 illustrates the ranked concepts in descending order of absolute weight across different runs. As shown in Figure 2f, validation set performance does not substantially improve beyond the top 25 concepts (median CI 0.69). We use the top 25 concept model’s test set performance for comparison with existing models in Figure 2a (hereafter referred to simply as ConSurv-Rule or C-Rule Top25).

In terms of median test performance, ConSurv models’ performance falls between XGBoost and DeepSurv. A Mann–Whitney U test confirms that ConSurv models, MLP, and DeepSurv perform significantly better than CPH. However, differences in performance among XGBoost, ConSurv models, MLP, and DeepSurv are not statistically significant (p-value *>* 0.05). The p-values for these comparisons are provided in Supplemental Table 2.

We identified three key advantages of ConSurv. First, it achieves performance comparable to existing models. Second, it demonstrates stable performance across runs, as evidenced by a CI standard deviation of 0.06 for ConSurv-XGB, lower than the 0.1 observed for XGBoost, indicating consistency comparable to black-box models. Finally, as a gray-box model, ConSurv prioritizes interpretability through concepts as a core feature, ensuring that predictions rely exclusively on these concepts, thereby enhancing fidelity (or faithfulness). In the following sections, we explore the interpretability offered through concepts by ConSurv.

### 2.4 Concepts demonstrate significant stratification between high- and low-risk groups

#### Interpretability of CPH

Among the existing models, the CPH model has the lowest performance but provides inherent interpretability, with feature coefficients directly reflecting their importance.

In Supplemental Figure 4, we present the top 40 features ranked by their absolute value of CPH coefficients. With these coefficients one can recognize the contribution made by every feature to the final prediction. These coefficients are faithful to the model as they are directly used for risk prediction.

#### Interpretability of DeepSurv (SHAP limitations)

In contrast, DeepSurv achieves the highest median performance among the evaluated models; however, interpreting its predictions requires post-hoc explanation techniques such as SHAP, which provides local explanations for individual patients. SHAP summary plots (Supplemental Figure 5), which aggregate feature importance across all patients, indicate that *days to birth* (age) is the most important feature driving Deep-Surv’s predictions. However, when we examined SHAP explanations for individual patients (Supplemental Figure 5), we observed substantial inconsistencies. For one patient, *days to birth* reduced the predicted risk (Supplemental Figure 6a), while for another, it increased the risk (Supplemental Figure 6b) and for a third patient (Supplemental Figure 6c), the feature had no effect at all (SHAP score of 0.0). Notably, for the third patient, where SHAP suggested age was irrelevant, changing only the age resulted in changes in the predicted risk (ΔLogRisk = 22 and 15 for ages 25 and 80, respectively). This example highlights a key limitation of SHAP: even for the most important features, the explanations provided may not reliably reflect how the model truly uses them. Similar inconsistencies were observed for other top feature *ajcc pathology t T* 2, which showed conflicting effects on risk across different patients. These findings underscore broader concerns regarding the faithfulness of post-hoc XAI techniques in accurately representing the inner workings of the models they are intended to explain, and there is growing research highlighting these limitations [11, 12, 66].

Our framework addresses these limitations through interpretable concepts that group features together, offering insight into potential interactions. Furthermore, because the final prediction relies exclusively on these concepts, they present faith-fulness in the predictions. Additionally, it provides concept importance through the learned weights assigned to each concept.

#### Interpretability of ConSurv

Original concept bottleneck models [20] use predefined, human-interpretable concepts with ground truth for supervised concept training. However, availability of such concepts in many cases is rare. This is especially true in biomedical applications where AI is employed to gather insights from the data. In our case, clinicians not only want predictions but also features important for those predictions. In such scenarios, unsupervised extraction of concepts is a possible strategy where the extracted concepts are later ratified with domain knowledge based on the features they capture. The work by [67] demonstrates a time-series concept bottleneck model using this strategy. We use a similar approach where we first extract concepts by grouping features using either XGBoost or RuleKit (See Figure 4b, details in Section 4.2). Supplemental Tables 4–9 provides the top five concepts for ConSurv-XGB and ConSurv-Rule.

We evaluate the effectiveness of each of the top five concepts from both models in distinguishing high- and low-risk groups using Kaplan-Meier survival plots. We obtained the output for all patients for each of the top five concepts using the ConSurv-XGB and ConSurv-Rule models. These outputs were then used to generate the KM plots presented in Figure 3 (c for ConSurv-XGB and d for ConSurv-Rule). For the ConSurv-XGB model, all concepts except concept 2 showed significant stratification (i.e., *p <* 0.05) between high- and low-risk patients. Similarly, for the ConSurv-Rule model, all concepts except concept 4 demonstrated significant differentiation (i.e., *p <* 0.05) between patient groups. This suggests that the top five most influential concepts in risk prediction from both models can effectively distinguish patients. In the next section, we analyze and interpret these top five concepts in the context of their biological relevance.

### 2.5 Concepts capture biological processes

The concepts used in ConSurv are inherently interpretable and are extracted through two distinct approaches: ConSurv-XGB, where concepts are extracted using XGBoost, and ConSurv-Rule, where RuleKit is employed to define the concepts. First we look at the interpretation of top 5 concepts from ConSurv-XGB. Following which we look at interpretation of top 5 concepts extracted from ConSurv-Rule.

#### Interpretation of top 5 ConSurv-XGB concepts

**Concept 1:** The top-ranked concept in ConSurv-XGB, which exhibits the most significant stratification between high- and low-risk groups. This concept comprises the features: age, lymph, IGLV2-23, and MIR1244-4. Age is a key factor in survival outcomes, with women under 40 and over 80 showing poorer prognosis [35]. Lymph node pathology is also crucial, as one pathway for breast cancer metastasis is through the lymphatic system and lymph nodes [68]. This is especially significant for women under 40 with axillary lymph node-negative breast cancer shown to have poor prognosis [35]. Older patients with a high lymph node ratio are shown to have a threefold increased risk of breast cancer death. [69, 70]. The IGLV family (here, IGLV2-23) is linked to older age at diagnosis and a distinct stromal microenvironment in breast cancer [71]. This concept connects age, lymph node pathology, and the IGLV gene. While we couldn’t find details of mechanism for MIR1244-2, there is increasing evidence of microRNAs’ role in breast cancer malignancy [72].

**Concept 2:** Comprises of NEAT1, TFF1, S100A6, RPS4Y1 and SETSIP genes. NEAT1 [73] promotes breast cancer, while TFF1 [74] and S100A6 [75] suppress it. All three interact with or affect estrogen expression [74, 76, 77]. RPS4Y1 [33], a male breast cancer marker, may serve as a proxy for estrogen status. SETSIP, while not directly linked, is associated with angiogenesis [78], potentially supporting tumor growth. This concept reflects estrogen response in breast cancer.

**Concept 3:** Includes four genes—AQP8, PCSK1, HSPB1, and LGALS4—along with one clinical feature, lymph node pathology. AQP8 is highly expressed in basal and luminal B breast cancer types, which have low estrogen receptor (ER) levels [79]. PCSK1 is upregulated in breast cancer, promoting tumor progression, estrogen dependency, and anti-estrogen resistance in cell lines [80]. HSPB1 is linked to metastasis via epithelial-to-mesenchymal transition and is correlated with lymph node status and estrogen receptors [81]. LGALS4 high expression is a good prognostic factor for LN-negative patients [82]. This concept highlights estrogen dependency (via AQP8, PCSK1, and HSPB1) and lymph node involvement (via lymph node pathology, HSPB1, and LGALS4).

**Concept 4:** Includes features COL11A2, TAGLN2, LPL, CPB1 and lymph node pathology. COL11A2 [83] and TAGLN2 [84, 85] are linked to lymph node pathology.

LPL promotes tumor growth by altering the microenvironment through lipid hydrolysis [86], while all three are potential therapeutic targets. CPB1 down-regulates tumor suppressors like SFRP1 and OS9 [87] and is up-regulated in lymph node-positive patients [88]. This concept highlights features associated with lymph node involvement and potential therapeutic targets.

**Concept 5:** Comprises of HBD, CD24, IGHV5-51, RPS4Y1 and age. The loss of the immunomodulatory HBD [89] and overexpression of CD24 [27, 90] promote metastasis by enabling escape from cell death. The IGHV family (e.g., IGHV5-51) is linked to vacuolation and degeneration in mouse breast tumorigenesis [91]. Together, these features suggest immune dysregulation, promoting cell death avoidance and tumor proliferation. However, the role of age and RPS4Y1 in conjunction with these genes requires further exploration.

We find ConSurv-XGB concepts are smaller in size (only 1–5 features per concept), with each capturing a biologically meaningful property. Further, understanding relationships between features within each concept often requires domain knowledge as seen for the five concepts discussed above.

#### Interpretation of top 5 ConSurv-Rule concepts

The concepts extracted from RuleKit tend to be larger (25–30 features per concept), leading us to hypothesize that they capture broader biological processes. To investigate this, we performed Overrepresentation Analysis (ORA) (See Section 4.8) to identify associated processes, with results listed in Table 3. We used the Hallmark [92] and C6 gene set databases in MSigD to determine relevant processes. Hallmark gene sets represent well-defined biological states with coherent expression patterns and were our primary reference. When no significant process were identified in Hallmark, we turned to C6. We found that all top 5 concepts in RuleKit can be associated with broader biological processes. Concept 1 primarily captures signaling pathways but also identifies hypoxia, a key feature of the tumor microenvironment that promotes angiogenesis in breast cancer [93]. Concept 2 is associated with immune-related processes, as indicated by the Hallmark allograft rejection and interferon-gamma response gene sets, both critical in breast cancer progression [94, 95]. Concept 4, which did not significantly differentiate between high- and low-risk patients (Figure 3d), lacks Hallmark features but captures the C6 oncogenic signature CTIP DN.V1 UP, linked to early-onset breast cancer [96].

**Table 3.**
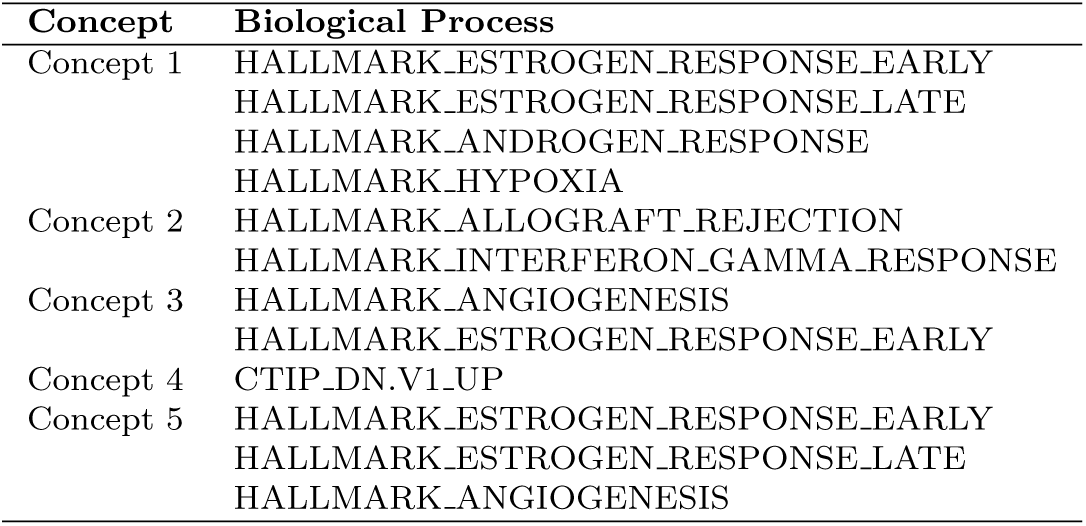
Biological processes of top 5 ConSurv-Rule concepts.

| Concept | Biological Process |
| --- | --- |
| Concept 1 | HALLMARK_ESTROGEN_RESPONSE_EARLY<br>HALLMARK_ESTROGEN_RESPONSE_LATE<br>HALLMARK_ANDROGEN_RESPONSE<br>HALLMARK_HYPOXIA |
| Concept 2 | HALLMARK_ALLOGRAFT_REJECTION<br>HALLMARK_INTERFERON_GAMMA_RESPONSE |
| Concept 3 | HALLMARK_ANGIOGENESIS<br>HALLMARK_ESTROGEN_RESPONSE_EARLY |
| Concept 4 | CTIP_DN.V1_UP |
| Concept 5 | HALLMARK_ESTROGEN_RESPONSE_EARLY<br>HALLMARK_ESTROGEN_RESPONSE_LATE<br>HALLMARK_ANGIOGENESIS |

RuleKit concepts exhibit two key characteristics: 1. Certain gene sets, such as estrogen response and angiogenesis, appears repeatedly across concepts (both processes are crucial for breast cancer survival [93, 97]); and 2. they often encompass multiple biological processes suggesting that RuleKit identifies high-level biological pattern.

Both ConSurv models provide biologically meaningful concepts. While ConSurv-XGB requires domain knowledge to interpret feature associations within each concept, ConSurv-Rule concepts can be more easily linked to well-established biological processes. Unlike conventional methods that rely on individual feature weights or costly and often unreliable post-hoc explanations, ConSurv models offer direct biological insights through their inherently interpretable design.

In the next section, we analyze quantitative properties of the concepts from both models.

### 2.6 Quantitative analysis of the interpretable concepts

In this section, we explore the quantitative aspects of the interpretable concepts in our framework. First, we conduct an exploratory analysis of the concepts (Figure 5) and then look at the stability of the concepts and robustness of the concept extraction.

**Fig. 5.**
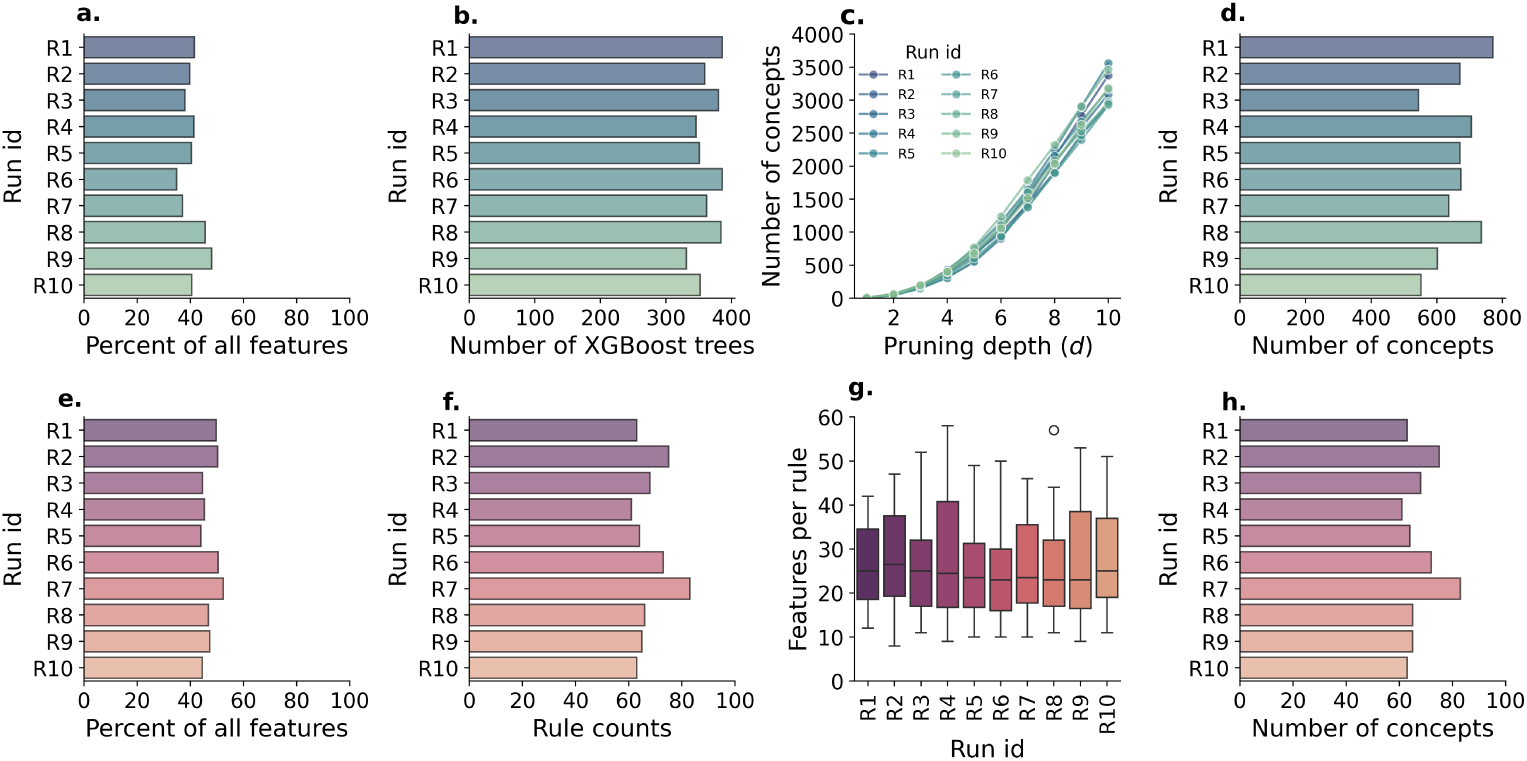
Exploratory analysis of XGBoost, RuleKit, and extracted concepts across ten runs. **a.** Trained XGBoost trees covered 35–50% of the 1,056 features in the dataset and generated 330–390 trees (**b**). **c.** The number of unique concepts extracted from trained trees drops significantly from 2,900–3,600 at depth *d* = 10 to 4–8 at *d* = 1. **d.** For trees with pruning depth *d* = 5 the number of unique concepts ranges from 540–770. **e.** RuleKit-generated rules cover 43–53% of the 1,056 available features. **f.** RuleKit produced 61–83 rules, with each rule containing an average of 25–30 conditions (**g**, Each box shows the 1st and 3rd quartiles, with the middle line indicating the median. Whiskers show the minimum and maximum, and outliers are shown as circles). **h.** Between 61–83 unique concepts were extracted, with one concept per rule.

Figure 5a shows that for each run, XGBoost utilizes between 35–50% of the total 1,056 features in the dataset. Each XGBoost run produces between 330–390 trees (Figure 5b). The depth of the tree dictates the number of unique concepts that can be extracted. As seen in Figure 5c, the number of concepts drops drastically from depth 10 to depth 1. Given that pruning depth 5 achieves performance comparable to depth 10, we extract concepts at this pruning depth, yielding approximately 540–770 unique concepts (Figure 5d).

Since the top 100 concepts contribute most to the predictive capability of ConSurv-XGB, we analyzed their origin. Specifically, we examined whether these concepts emerge from early or late iterations of XGBoost training. Figure 6a demonstrates that most highly ranked concepts emerge during the initial and final iterations, a trend consistent across runs. Figure 6b shows that larger-sized concepts (based on number of features within each concept) are more frequent in the top 100 concepts, with concepts of size 5 being predominant.

**Fig. 6.**
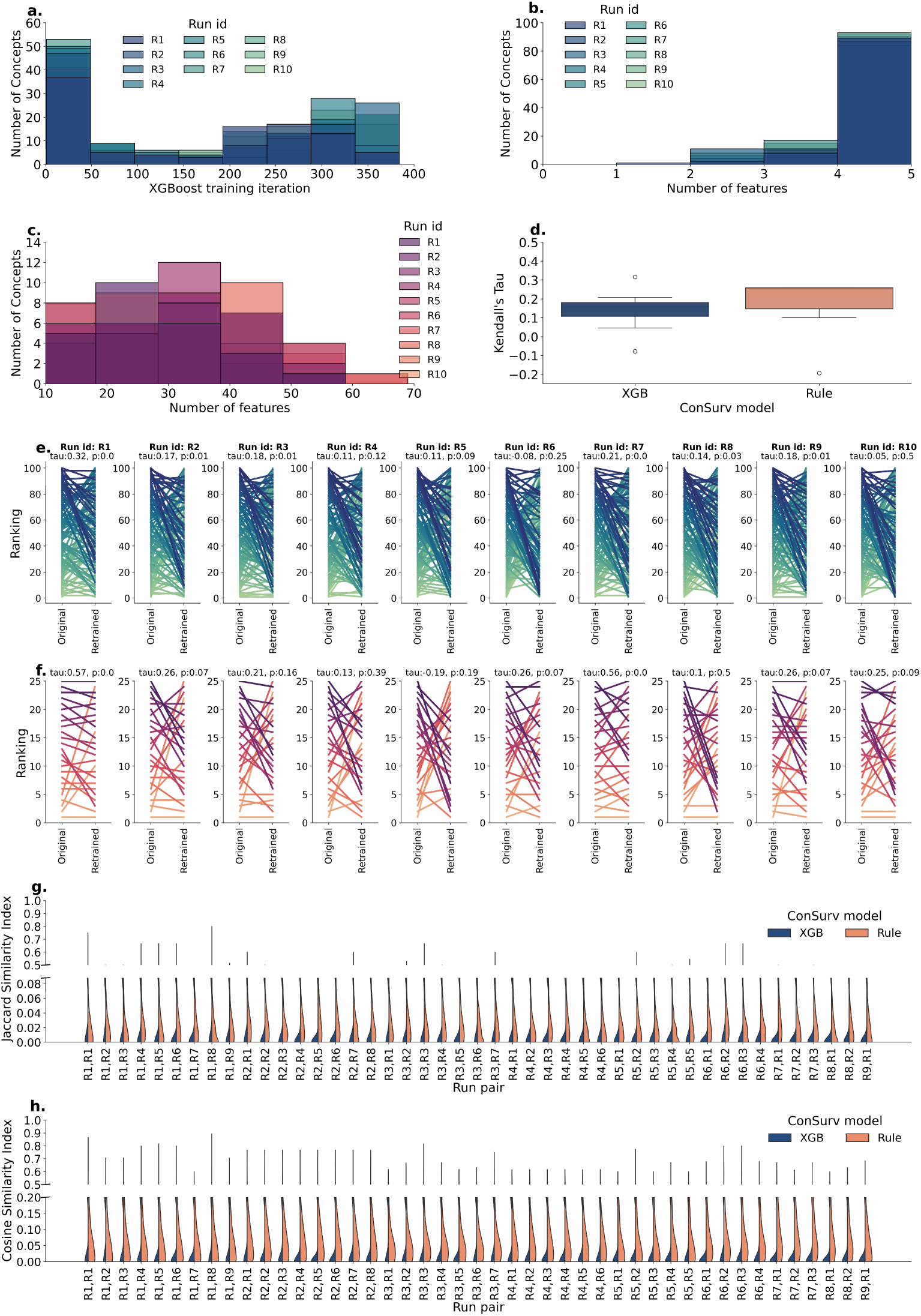
Quantitative analysis of extracted concepts across ten runs. **a.** The distribution of the top 100 concepts from ConSurv-XGB shows that most originate from XGBoost trees generated in early or late training iterations. **b.** Among the top 100 concepts from ConSurv-XGB, the majority comprise 5 features, with gradually fewer concepts having 4, 3, or 2 features. **c.** For ConSurv-Rule, the top 25 concepts typically contain 20–40 features per concept. **d.** To assess the stability of concept rankings, the rankings of concepts in ConSurv-XGB and ConSurv-Rule were compared to their original rankings in the ConSurv-XGB-all and ConSurv-Rule-all models, respectively, using Kendall’s Tau. Kendall’s Tau values for ConSurv-XGB ranged from -0.8 to_1_0_7_.32, indicating minimal similarity, while ConSurv-Rule concepts showed higher similarity with Kendall’s Tau values between 0.13 and 0.57. Each box shows the 1st and 3rd quartiles, with the middle line indicating the median. Whiskers show the minimum and maximum, and outliers are shown as circles. **e.** Kendall’s Tau values across ten runs showing change in the rankings of the top 100 ConSurv-XGB concepts between the original ranking and the new ranking. **f.** Kendall’s Tau values across ten runs reflecting changes in the rankings of the top 25 ConSurv-Rule concepts between the original ranking and the new ranking. **g.** The distribution of Jaccard Similarity Index between concepts captured across run pairs shows very limited overlap for both ConSurv-XGB and ConSurv-Rule concepts. **h.** Similarly, the distribution of Cosine Similarity Index indicates limited overlap between concepts across runs for both methods. However, ConSurv-Rule concepts exhibit greater overlap than ConSurv-XGB concepts on both similarity indices.

In each run of RuleKit, between 43–53% of the 1,056 dataset features were utilized that produced 61–83 rules, with each rule containing an average of 25–30 conditions (Figure 5e–g). Unlike XGBoost, each unique rule roughly translates to one unique concept, resulting in 61–83 unique concepts per run (Figure 5h).

#### Stability

Next, we examined whether training the model using only the top-ranking concepts impacts their rankings. A model that preserves most of the concept rankings would suggest that these concepts and their importance remain stable. We compared the ranking of 100 concepts in ConSurv-XGB model with original rankings from ConSurv-XGB with all concepts. We used Kendall’s Tau to assess the similarity of rankings and found that it ranged between 0.08 to 0.31 (plotted in Figure 6d,e) which indicates there is little to no correlations between the two rankings. Similarly, we compared the ranking of the 25 concepts in ConSurv-Rule model with original rankings from ConSurv-Rule with all concepts. We found that the Kendall’s Tau value ranged between 0.02 to 0.56 (plotted in Figure 6d, f) which indicates there is little correlations (better than ConSurv-XGB) between the two rankings. These results, which show a high mismatch between the original and retrained rankings for both ConSurv-XGB and ConSurv-Rule concepts, indicate that the concepts and their importance are not stable (i.e. the concepts are not stable).

#### Robustness

The concept extraction process can be considered robust if the concepts extracted across multiple runs exhibit a high degree of similarity. We evaluated the similarity of concepts across runs in both ConSurv models. Concepts represent sets of features, with sizes ranging from 1–5 features for ConSurv-XGB and averaging 25–30 features for ConSurv-Rule. To measure similarity, we used the commonly applied Jaccard Similarity Index (SI). However, as Jaccard SI is influenced by set size, we also used the Cosine SI, which focuses on overall similarity between sets. Figures 6g,h shows the distributions of Jaccard and Cosine SI across the 45 run pairs (10 runs yields 45 pairs), comparing every concept from one run to every concept in another run. For each pair, comparing 100 concepts in ConSurv-XGB results in 4,950 comparisons and 300 comparisons for the 25 concepts in ConSurv-Rule. ConSurv-Rule concepts were noticeably more consistent across runs compared to those from XGBoost, as indicated by higher Jaccard and Cosine similarity indices. This consistency is further supported by two observations: First, a trained RuleKit model utilizes more features per run than a trained XGBoost model (43%–53% vs. 35%–50%, respectively), allowing for more potential overlap across runs. Second, 44 features were consistently present across RuleKit runs, compared to only 14 features in XGBoost runs. However, despite this relative difference, the overall similarity indices for both remained quite low, indicating limited overlap in the concepts across runs.

In summary, ConSurv-XGB concepts are noticeably smaller (1–5 features per concept) than ConSurv-Rule concepts (25–30 features per concept). Similarity in concepts across runs is limited for both models. However, in contrast to ConSurv-XGB, ConSurv-Rule concepts are more consistent across runs. Additionally, concepts from both model appear not stable, as their rankings change after retraining.

## 3 Discussion and Conclusion

Our study aims to improve interpretability in survival analysis while balancing the trade-off with performance. Through an evaluation of existing models and our proposed ConSurv framework, we demonstrated that incorporating interpretable concepts enhances model transparency without compromising performance.

### Trade-off between interpretability and accuracy

The CPH, a fully white-box approach, offers inherent interpretability via feature coefficients but suffers from the lowest median performance. Conversely, black-box models such as DeepSurv achieve the highest median performance but require unreliable posthoc explanations. ConSurv, a gray-box model, successfully balances these concerns. By leveraging interpretable concepts from ConSurv-XGB and ConSurv-Rule, we achieve performance similar to high performing models while providing human-interpretable insights. Importantly, ConSurv-XGB demonstrates lower CI variance than XGBoost, indicating increased stability of the performance.

### Concepts enable significant stratification among patients

A key advantage of our framework lies in its ability to extract and leverage meaningful concepts in survival prediction. Unlike post-hoc methods that provide individual feature importance without capturing interactions, ConSurv integrates concepts directly into the model. This allows a more interpretable and faithful predictions with biologically relevant representation of survival risk factors. This is evident in our analysis of concepts and their biological significance. Additionally, Kaplan-Meier survival plots (Figure 3) show that top ConSurv-XGB and ConSurv-Rule provides significant survival stratification between high- and low-risk patients.

### Biological relevance of concepts

To assess the interpretability and biological relevance of our learned concepts, we performed Over-Representation Analysis (ORA) to visualize gene interaction networks. The extracted concepts align with known biological processes relevant to cancer survival, such as hypoxia, angiogenesis, etc. We apply very stringent condition that only Hallmark and C6 databases were used (with preference for Hallmark) to ensure only most relevant processes are obtained. However, this can be left at the discretion of the end user to set the criteria.

### Stability and reproducibility of extracted concepts

An important consideration in modeling approach is the consistency of learned concepts across different runs. Our analysis of concept robustness (Figure 6) reveals that ConSurv-Rule concepts exhibit greater consistency across runs compared to ConSurv-XGB concepts. However, despite this advantage, overall concept similarity remains low, indicating variability in the specific features grouped within each concept. Additionally, the stability of concepts for both models was low. More research is needed to develop strategies for extracting concepts more effectively, ensuring greater consistency across different runs. Reducing variability in concept formation across random data splits will be crucial for enhancing both the stability and robustness of learned concepts, thereby allowing the ConSurv framework to be more readily used.

### Limitations

Despite the advantages of ConSurv, some limitations warrant further investigation. First, the variability in concept extraction across runs suggests a need for more robust feature selection or concept aggregation strategies. One potential approach is to integrate ensemble learning techniques that uniformize concept extraction across multiple runs. Additionally, while ConSurv-Rule concepts are relatively robust, they include many features per concept which capture multiple processes, and exhibit redundancy across concepts. Future work could explore hybrid approaches that optimize both concept size and redundancy. With only 1–5 features per concept, ConSurv-XGB concepts are small, but interpreting their biological meaning requires extensive domain knowledge.

## Conclusion

In conclusion, our study introduces ConSurv, a gray-box survival analysis framework that balances interpretability and accuracy by leveraging concept-based learning. Through comparisons with existing models, we demonstrate that ConSurv achieves high predictive performance while maintaining transparency in its decision-making with biologically relevant concepts. This unique ability positions ConSurv as a valuable tool for clinical decision support, enabling both accurate risk prediction and actionable insights. Future research will focus on robust concept extraction and exploring broader applications in precision medicine.

## 4 Methods

### 4.1 Contrast Set Mining

Association rule mining is used in problems with multi-dimensional datasets to identify relations between features [98]. These associations takes a form of “if-then” rules. Contrast set mining [99–101] is a special type of association rule mining that identifies key differences between groups within a dataset. It specifically aims to find attribute-value (i.e. feature and its value) combinations, known as contrast sets, that vary significantly in frequency or magnitude across these groups.

For a dataset *D* containing two classes—positive (*P*) and negative (*N*)—a contrast set *S* is defined as a combination of attribute-value pairs (e.g., *A*1 = *v*1 *∧ A*2 = *v*2, where *v*1 and *v*2 are specific values of the attributes). The support of *S* within positive or negative class, represented as support(*S, P*) or support(*S, N*), is the proportion of instances in class that satisfy *S*. A contrast set *S* is deemed significant if there is at least one pair of classes, *P* and *N*, for which the support difference support(*S, P*) support(*S, N*) exceeds a given threshold *δ* and is statistically significant. For multiclass problems, it employs a one-vs-all strategy to generate rules specific to each class.

In this work, we leverage the RuleKit [101, 102] package to extract contrast sets.

### 4.2 ConSurv

ConSurv is an interpretable concept bottleneck model that relies solely on concepts for its final predictions. Its interpretability stems from the use of interpretable concepts, while maintaining complete faithfulness by basing predictions exclusively on a linear combination of these concepts, as shown below:

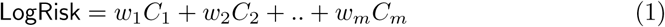

where *C*_1_..*C_m_* are *m* interpretable concepts and *w*_1_..*w_m_* their respective weights. ConSurv has two layers: one hidden layer and one *m*-sized concept layer. The architecture of ConSurv is designed such that each concept receives input only from the features grouped for that concept (see Figure 4a). The concept layer output is normalized to values between 0–1 to ensure that the impact of each concept is influenced only by the static weight *w* for that concept.

To extract concepts, we rely on trained XGBoost trees and RuleKit rules, as illustrated in Figure 4b. Contrast set rule are a form of association rule learning which find patterns among the features that aim to distinguish between different groups. As illustrated in Figure 4, for each learned rule, the features are collected and grouped into concepts. On the other hands, tree based algorithms such as XGBoost are good at identifying features which are non-linearly related to each other and output. For XGBoost, concepts are extracted from each depth level of trained tree. In the illustration of Figure 4, the tree has two levels nodes, *Node Pathology* feature at root and *Beta hemoglobin* feature at level 1. There the extracted concept from root level is just the *Node Pathology* feature (Concept 1) and from level 1 is *Node Pathology* and *Beta − hemoglobin* feature (Concept 2).

### 4.3 Hyperparameter tuning and model training

To ensure optimal parameters for model training, we conducted hyperparameter tuning for CPH, XGBoost, RuleKit, DeepSurv, MLP, and ConSurv. Ten runs were performed, with data splits generated using different random seeds (run IDs and their corresponding seeds are listed in Supplemental Table 1). For each run, the data was split into 70% training, 15% validation, and 15% test sets. Hyperparameter tuning was performed on the training and validation sets across all ten runs. The test sets were used to evaluate performance of the tuned models. The categorical features were one hot encoded and continuous features were min-max normalized.

The parameter combinations evaluated for each model are detailed in Supplemental Table 3, with the best-performing values highlighted in the last column. For ConSurv, the optimal parameters from MLP were adopted, with only the learning rate tuned separately. For XGBoost, DeepSurv, MLP, and ConSurv models, early stopping was applied with a patience of 50 rounds. Adaptive Moment Estimation (Adam) [103] was used for the gradient descent algorithm. The same training set used for training the ConSurv model was also used to train the corresponding XGBoost and RuleKit models for concept extraction.

### 4.4 Concordance Index

Concordance Index or CI [104, 105] is a used widely metric for evaluating performance of survival models. It is calculated as:

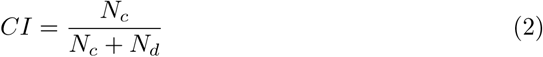

Where, *N_c_* is number of concordant pairs such that for patients *i* and *j*, *NN* (*x_i_*) *< NN* (*x_j_*), *T_i_ > T_j_* and *e_j_* = 1. *N_d_* is number of discordant pairs such that for patients *i* and *j*, *NN* (*x_i_*) *> NN* (*x_j_*), *T_i_ > T_j_* and *e_j_* = 1. *NN* (.) is predicted LogRisk, *T* is time-to-event and *e* is event variable.

### 4.5 Set similarity indices

Jaccard similarity index for set *A* and *B* is given by Equation (3). It ranges from 0–1 with 0 indicating no overlap or similarity and 1 indicating perfect overlap or similarity.

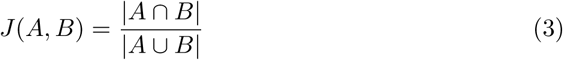

Cosine similarity index for set *A* and *B* is given by Equation (4). First the features from the two concepts being compared are one hot encoded into vectors *a* and *b* and then the Cosine similarity is calculated. The range of Cosine similarity is from -1–1, however, in our case of binary vectors, it ranges only from 0–1.

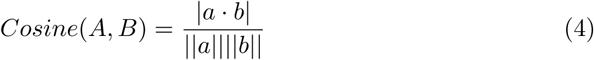

### 4.6 Objective function: Cox-loss

Deep learning models for survival analysis such as AutoSurv and DeepSurv often employ Cox-loss for training the models. We used the same loss function in training ConSurv and MLP models. The loss function is as follows (from [4, 8]):

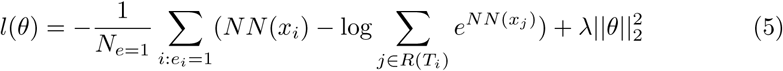

where *θ* are model parameters, *NN* (*xi*) (LogRisk) is model output for input *xi*, *λ* is *l*2 regularization parameter, *e* is the event variable, *Ne*=1 are number of patients with an observable event and *R*(*Ti*) are patients still at risk of failure at time *t*.

### 4.7 Gene Set Variation Analysis

Gene Set Variation Analysis (GSVA) is a non-parametric, unsupervised method that estimates the variation of pathway activity (or gene set enrichment) across all samples in a transcriptomic dataset [21]. Rather than examining individual gene expression, GSVA computes an enrichment score per sample for each predefined gene set, effectively transforming the gene-expression matrix into a sample-by-gene-set score matrix. In our work, we used GSVA to aggregate the genes associated with each concept (rule) into a single “concept activity score” per patient. The aggregated GSVA scores enable stratification of patients into survival groups, which we visualize using Kaplan-Meier plots.

### 4.8 Over-Representation Analysis

We performed Over-Representation Analysis (ORA) to identify biological pathways or gene sets enriched among top genes from each ConSurv rule. Using the Molecular Signatures Database (MSigDB) collections [92] (H - Hallmark gene sets and C6 - oncogenic signatures) and all measured genes as a background set, we applied Fisher’s exact test to evaluate whether a given genes in a concept appeared more frequently in any particular gene set than expected by chance. We corrected for multiple testing using the Benjamini–Hochberg method (FDR *<* 0.05). These enriched pathways provide functional context for each rule, underscoring potential mechanisms underlying risk stratification.

## Data Availability

The breast cancer dataset used in this study is available at https://anonymous.4open.science/r/ConSurv-13BE.

## Code availability

All models were implemented using Python-3.9. Following packages were used: xgboost (for XGBoost model), RuleKit (for Contrast Set Rule mining), lifelines (for CPH and concordance index), TorchSurv [106] (for loss function), PyTorch (For deep learning models - DeepSurv, MLP and ConSurv). The code and complete list of packages used is available at: https://anonymous.4open.science/r/ConSurv-13BE

## Acknowledgment

This work and authors P.B., A.R., are supported by the European Union’s Horizon 2020 research and innovation programme under grant agreement no. 101017453. A.V. is supported by the “UNREAL: Unified Reasoning Layer for Trustworthy ML” project (EP/Y023838/1) selected by the ERC and funded by UKRI EPSRC. This work has made use of the resources provided by the Edinburgh Compute and Data Facility (ECDF) (http://www.ecdf.ed.ac.uk/).

## Author contributions

P.B ., A.V., and A.R., conceptualized and designed experiments. P.B. and T.W. conducted the experiments and generated results. A.R. supervised the project. P.B., T.W., A.V., A.R. contributed to writing the manuscript.

