## Supplemental Material for "Can interpretability and accuracy coexist in cancer survival analysis?"

Table 1: Run id with corresponding seed

| Run id | Seed |
|---|---|
| R1 | 0 |
| R2 | 7 |
| R3 | 42 |
| R4 | 200 |
| R5 | 777 |
| R6 | 999 |
| R7 | 1303 |
| R8 | 1995 |
| R9 | 1996 |
| R10 | 2405 |

Table 2: Mann–Whitney U test results on test CI distribution

| **model** | **model** | **statistic** | **p-value** |
|---|---|---|---|
| C-Rule Top25 | C-XGB Top100 | 46.0 | 0.791337 |
| C-Rule Top25 | CPH | 78.0 | **0.037635** |
| C-Rule Top25 | DeepSurv | 36.0 | 0.307489 |
| C-Rule Top25 | MLP | 37.0 | 0.344704 |
| C-Rule Top25 | XGBoost | 47.0 | 0.850107 |
| C-XGB Top100 | CPH | 84.0 | **0.011330** |
| C-XGB Top100 | DeepSurv | 42.0 | 0.570750 |
| C-XGB Top100 | MLP | 42.0 | 0.570750 |
| C-XGB Top100 | XGBoost | 55.0 | 0.733730 |
| CPH | DeepSurv | 17.0 | **0.014019** |
| CPH | MLP | 16.0 | **0.011330** |
| CPH | XGBoost | 31.0 | 0.161972 |
| DeepSurv | MLP | 54.0 | 0.791337 |
| DeepSurv | XGBoost | 60.0 | 0.472676 |
| MLP | XGBoost | 58.0 | 0.570750 |

Table 3: Parameter list for hyperparameter tuning

| Parameter | Values | Best value |
|---|---|---|
| | **CPH** | |
| *penalizer* | $[0.01, 1]$ | 1 |
| *l1_ratio* | $[0.5, 1]$ | 0.5 |
| | **XGBoost** | |
| *max_depth* | $[5, 10, 20, 40]$ | 10 |
| *learning_rate* | $[10^{-1}, 10^{-2}, 10^{-3}]$ | $10^{-2}$ |
| *n_estimator* | $[10, 50, 100, 250, 500, 750, 1000, 2000]$ | 500 |
| *lambda* | $[10^{-1}, 10^{-2}, 10^{-3}]$ | $10^{-2}$ |
| *alpha* | $[10^{-1}, 10^{-2}]$ | $10^{-1}$ |
| *aft_loss_distribution* | $[0.1, 0.5, 1.2]$ | 1.2 |
| | **RuleKit** | |
| *measures* | $[c2, rss, correlation]$ | *rss* |
| *minsupp_new* | $[3, 5, 7, 9, 11, 13]$ | 3 |
| | **DeepSurv** | |
| *lr* | $[10^{-1}, 10^{-3}, 10^{-5}]$ | $10^{-5}$ |
| *hidden_size* | $[64, 128, 256]$ | 128 |
| *l2_reg* | $[0.1, 0.01, 0.001]$ | 0.01 |
| *batch_size* | $[32, 64]$ | 64 |
| *dropout* | $[0, 0.1, 0.2]$ | 0.2 |
| | **MLP** | |
| *batch_size* | $[16, 32, 64]$ | 64 |
| *L2_reg* | $[10^{-1}, 10^{-2}, 10^{-3}]$ | $10^{-3}$ |
| *learning_rate* | $[10^{-1}, 10^{-3}, 10^{-5}]$ | $10^{-5}$ |
| *hidden_size* | $[32, 64, 128, 256, 512]$ | 64 |
| *n_layer* | $[1, 2, 3]$ | 2 |
| | **ConSurv** | |
| *learning_rate* | $[10^{-1}, 10^{-3}, 10^{-5}]$ | $10^{-3}$ |

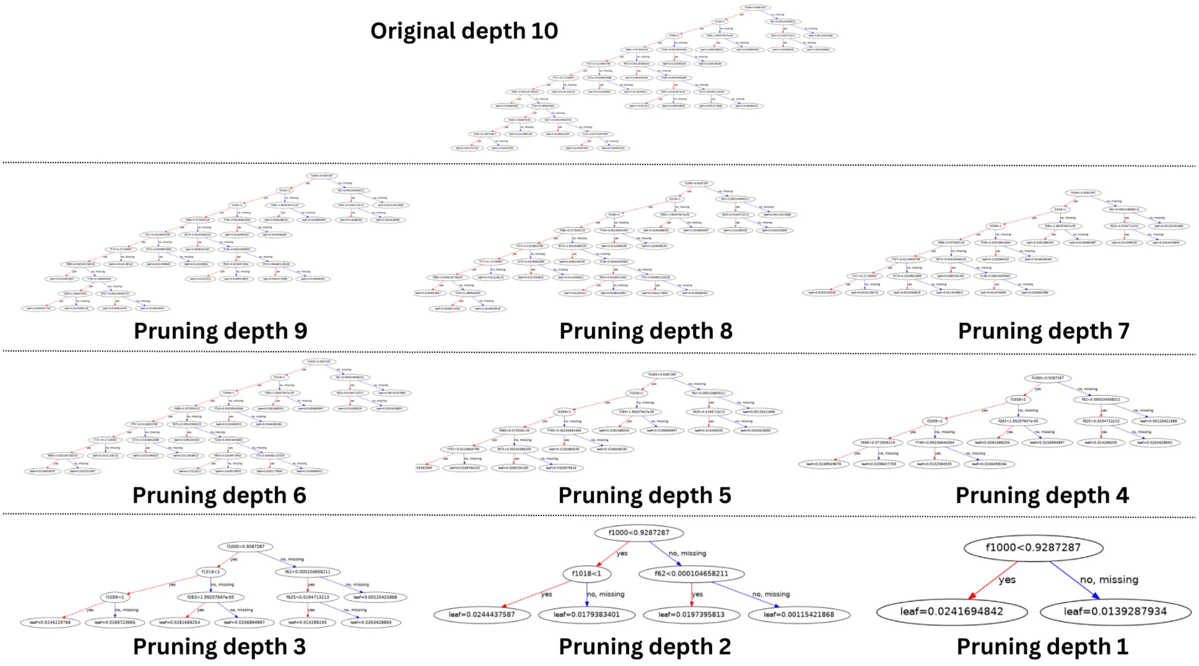


Figure 1: Pruning an XGBoost tree from original depth of 10 to a pruning depth of 1

Table 4: ConSurv-XGB top 5 concepts for Run 6

| Features | Alternative Name |
|---|---|
| | **Concept 1, Rank 1** |
| demographic.days_to_birth | Age |
| ENSG00000283498.1 | MIR1244-2 |
| ENSG00000211660.3 | IGLV2-23 |
| ajcc_pathologic_n_N0 (i-) | Lymph node pathology |
| | **Concept 2, Rank 2** |
| ENSG00000245532.9 | NEAT1 |
| ENSG00000160182.3 | TFF1 |
| ENSG00000230667.6 | SETSIP |
| ENSG00000197956.10 | S100A6 |
| ENSG00000129824.16 | RPS4Y1 |
| | **Concept 3, Rank 3** |
| ENSG00000103375.11 | AQP8 |
| ENSG00000175426.11 | PCSK1 |
| ENSG00000106211.10 | HSPB1 |
| ENSG00000171747.9 | LGALS4 |
| ajcc_pathologic_n_N2 | Lymph node pathology |
| | **Concept 4, Rank 4** |
| ENSG00000175445.17 | LPL |
| ENSG00000204248.11 | COL11A2 |
| ENSG00000153002.12 | CPB1 |
| ENSG00000158710.15 | TAGLN2 |
| ajcc_pathologic_n_N1b | Lymph node pathology |
| | **Concept 5, Rank 5** |
| ENSG00000211966.2 | IGHV5-51 |
| ENSG00000129824.16 | RPS4Y1 |
| ENSG00000223609.11 | HBD |
| ENSG00000185275.6 | CD24P4 |
| days_to_birth | Age |

Table 5: ConSurv-Rule Concept 1, Rank 1 for Run 6

| **Features** | **Alternative Name** |
|---|---|
| ENSG00000197249.14 | SERPINA1 |
| ENSG00000146648.19 | EGFR |
| ENSG00000141424.13 | SLC39A6 |
| ENSG00000170345.10 | FOS |
| ENSG00000225972.1 | MTND1P23 |
| ENSG00000189058.9 | APOD |
| ENSG00000172551.11 | MUCL1 |
| ENSG00000134827.8 | TCN1 |
| ENSG00000211639.2 | IGLV4-60 |
| ENSG00000141750.7 | STAC2 |
| ENSG00000134258.17 | VTCN1 |
| ENSG00000122711.9 | SPINK4 |
| ENSG00000170421.12 | KRT8 |
| ENSG00000160182.3 | TFF1 |
| ENSG00000147256.12 | ARHGAP36 |
| ENSG00000054938.16 | CHRDL2 |
| ENSG00000211637.2 | IGLV4-69 |
| ENSG00000204434.6 | POTEKP |
| ENSG00000263934.5 | SNORD3A |
| ENSG00000253818.1 | IGLV1-41 |
| ENSG00000138696.11 | BMPR1B |
| ENSG00000263639.7 | MSMB |
| ENSG00000167754.13 | KLK5 |
| ENSG00000106541.12 | AGR2 |
| ENSG00000159335.18 | PTMS |
| ENSG00000198938.2 | MT-CO3 |
| ENSG00000197253.13 | TPSB2 |
| ENSG00000139329.5 | LUM |
| ENSG00000134201.12 | GSTM5 |
| ENSG00000163736.4 | PPBP |
| ENSG00000211660.3 | IGLV2-23 |
| ENSG00000012660.14 | ELOVL5 |
| ENSG00000123999.5 | INHA |
| ENSG00000253802.2 | AC105999.2 |
| ENSG00000146678.10 | IGFBP1 |
| ENSG00000164120.14 | HPGD |
| ENSG00000113739.10 | STC2 |
| ENSG00000094755.17 | GABRP |
| ENSG00000200087.1 | SNORA73B |
| ENSG00000112306.8 | RPS12 |
| ENSG00000167258.15 | CDK12 |
| ENSG00000178795.9 | GDPD4 |
| ENSG00000286775.1 | - |
| ENSG00000164756.12 | SLC30A8 |
| ENSG00000171747.9 | LGALS4 |
| ENSG00000222880.1 | RN7SKP261 |
| ENSG00000109321.11 | AREG |
| ENSG00000211655.3 | IGLV1-36 |
| ENSG00000163631.17 | ALB |
| ENSG00000141232.5 | TOB1 |
| ENSG00000143125.6 | PROK1 |
| ENSG00000108679.13 | LGALS3BP |
| ENSG00000172724.12 | CCL19 |
| ENSG00000173335.5 | CST9 |
| ENSG00000187653.11 | TMSB4XP8 |
| ENSG00000179593.16 | ALOX15B |
| ENSG00000211950.2 | IGHV1-24 |
| ENSG00000140988.16 | RPS2 |
| ajcc_pathologic_stage I | Pathology stage |
| ajcc_pathologic_stage IA | Pathology stage |
| ajcc_pathologic_stage IB | Pathology stage |
| ajcc_pathologic_stage II | Pathology stage |
| ajcc_pathologic_stage IIA | Pathology stage |
| ajcc_pathologic_stage IIB | Pathology stage |
| ajcc_pathologic_stage III | Pathology stage |
| ajcc_pathologic_stage IIIA | Pathology stage |
| ajcc_pathologic_stage IIIB | Pathology stage |
| ajcc_pathologic_stage IIIC | Pathology stage |
| ajcc_pathologic_stage IV | Pathology stage |
| ajcc_pathologic_stage X | Pathology stage |

Table 6: ConSurv-Rule Concept 2, Rank 2 for Run 6

| **Features** | **Alternative Name** |
|---|---|
| ENSG00000167768.4 | KRT1 |
| ENSG00000128016.7 | ZFP36 |
| ENSG00000105894.12 | PTN |
| ENSG00000170893.4 | TRH |
| ENSG00000200164.1 | RF00019 |
| ENSG00000140986.8 | RPL3L |
| ENSG00000218175.2 | AC016739.1 |
| ENSG00000105388.16 | CEACAM5 |
| ENSG00000199629.1 | RNU1-14P |
| ENSG00000173821.19 | RNF213 |
| ENSG00000238034.1 | AL109807.1 |
| ENSG00000241351.3 | IGKV3-11 |
| ENSG00000165949.12 | IFI27 |
| ENSG00000137673.9 | MMP7 |
| ENSG00000109072.14 | VTN |
| ENSG00000147604.14 | RPL7 |
| ENSG00000161634.12 | DCD |
| ENSG00000099194.6 | SCD |
| ENSG00000286339.1 | - |
| ENSG00000207205.1 | RNVU1-15 |
| ENSG00000115414.21 | FN1 |
| ENSG00000212283.1 | SNORD89 |
| ENSG00000171246.6 | NPTX1 |
| ENSG00000198888.2 | MT-ND1 |
| ENSG00000211866.1 | TRAJ23 |
| ENSG00000204936.10 | CD177 |
| ENSG00000229344.1 | MTCO2P12 |
| ENSG00000104267.10 | CA2 |
| ENSG00000163220.11 | S100A9 |
| ENSG00000121769.8 | FABP3 |
| ENSG00000229859.10 | PGA3 |
| ENSG00000240409.1 | MTATP8P1 |
| ENSG00000133048.13 | CHI3L1 |
| ENSG00000211598.2 | IGKV4-1 |
| ENSG00000281990.1 | IGHV1-69-2 |
| ENSG00000206503.13 | HLA-A |
| ENSG00000212605.1 | RNU1-56P |
| ENSG00000273716.2 | AC092670.1 |
| ENSG00000166710.21 | B2M |
| ENSG00000096006.12 | CRISP3 |
| ENSG00000075388.4 | FGF4 |
| ENSG00000242265.6 | PEG10 |
| ENSG00000161055.4 | SCGB3A1 |
| ENSG00000162267.12 | ITIH3 |
| ENSG00000240216.7 | CPHL1P |
| ENSG00000151224.13 | MAT1A |

Table 7: ConSurv-Rule Concept 3, Rank 3 for Run 6

| **Features** | **Alternative Name** |
|---|---|
| ENSG00000171428.15 | NAT1 |
| ENSG00000170345.10 | FOS |
| ENSG00000178473.7 | UCN3 |
| ENSG00000136881.12 | BAAT |
| ENSG00000182853.12 | VMO1 |
| ENSG00000189058.9 | APOD |
| ENSG00000141736.14 | ERBB2 |
| ENSG00000170807.12 | LMOD2 |
| ENSG00000175899.15 | A2M |
| ENSG00000183666.17 | GUSBP1 |
| ENSG00000129988.6 | LBP |
| ENSG00000116882.15 | HAO2 |
| ENSG00000141750.7 | STAC2 |
| ENSG00000132703.4 | APCS |
| ENSG00000278233.1 | RNA5-8SN3 |
| ENSG00000054938.16 | CHRDL2 |
| ENSG00000156885.6 | COX6A2 |
| ENSG00000124107.5 | SLPI |
| ENSG00000133110.15 | POSTN |
| ENSG00000212907.2 | MT-ND4L |
| ENSG00000159388.6 | BTG2 |
| ENSG00000248746.6 | ACTN3 |
| ENSG00000265929.1 | MIR5195 |
| ENSG00000263639.7 | MSMB |
| ENSG00000118849.10 | RARRES1 |
| ENSG00000228253.1 | MT-ATP8 |
| ENSG00000174697.5 | LEP |
| ENSG00000211976.2 | IGHV3-73 |
| ENSG00000016490.16 | CLCA1 |
| ENSG00000197253.13 | TPSB2 |
| ENSG00000142089.16 | IFITM3 |
| ENSG00000248144.6 | ADH1C |
| ENSG00000143248.13 | RGS5 |
| ENSG00000005381.8 | MPO |
| ENSG00000249780.1 | AC093809.1 |
| ENSG00000012660.14 | ELOVL5 |
| ENSG00000248779.1 | AC093297.1 |
| ENSG00000141753.7 | IGFBP4 |
| ENSG00000253755.1 | IGHGP |
| ENSG00000186009.5 | ATP4B |
| ENSG00000183607.10 | GKN2 |
| ENSG00000109072.14 | VTN |
| ENSG00000261409.1 | AL035425.3 |
| ENSG00000158104.11 | HPD |
| ENSG00000124939.6 | SCGB2A1 |
| ENSG00000225630.1 | MTND2P28 |
| ENSG00000102837.7 | OLFM4 |
| ENSG00000131771.14 | PPP1R1B |
| ENSG00000164128.7 | NPY1R |
| ENSG00000101443.18 | WFDC2 |
| ENSG00000140459.18 | CYP11A1 |
| ENSG00000172724.12 | CCL19 |
| ENSG00000179593.16 | ALOX15B |
| ENSG00000159167.12 | STC1 |
| days_to_birth | Age |

Table 8: ConSurv-Rule Concept 4, Rank 4 for Run 6

| **Features** | **Alternative Name** |
|---|---|
| ENSG00000125999.11 | BPIFB1 |
| ENSG00000166426.8 | CRABP1 |
| ENSG00000211668.2 | IGLV2-11 |
| ENSG00000104368.19 | PLAT |
| ENSG00000092054.13 | MYH7 |
| ENSG00000164326.5 | CARTPT |
| ENSG00000143632.14 | ACTA1 |
| ENSG00000019582.15 | CD74 |
| ENSG00000205361.8 | MT1DP |
| ENSG00000229314.5 | ORM1 |
| ENSG00000239855.1 | IGKV1-6 |
| ENSG00000240386.3 | LCE1F |
| ENSG00000248527.1 | MTATP6P1 |
| ENSG00000170367.5 | CST5 |
| ENSG00000283907.1 | AD000090.1 |
| ENSG00000230715.3 | AC018638.2 |
| ENSG00000096384.20 | HSP90AB1 |
| ENSG00000241755.1 | IGKV1-9 |
| ENSG00000171246.6 | NPTX1 |
| ENSG00000204936.10 | CD177 |
| ENSG00000171345.13 | KRT19 |
| ENSG00000145824.13 | CXCL14 |
| ENSG00000124107.5 | SLPI |
| ENSG00000167676.4 | PLIN4 |
| ENSG00000169347.17 | GP2 |
| ENSG00000141744.4 | PNMT |
| ENSG00000221421.1 | MIR1283-1 |
| ENSG00000231414.1 | AC016700.2 |
| ENSG00000118271.12 | TTR |
| ENSG00000244461.1 | LINC02077 |
| ENSG00000132693.12 | CRP |
| ENSG00000167531.6 | LALBA |
| ajcc_pathologic_n_N0 | Lymph node pathology |
| ajcc_pathologic_n_N0 (i+) | Lymph node pathology |
| ajcc_pathologic_n_N0 (i-) | Lymph node pathology |
| ajcc_pathologic_n_N0 (mol+) | Lymph node pathology |
| ajcc_pathologic_n_N1 | Lymph node pathology |
| ajcc_pathologic_n_N1a | Lymph node pathology |
| ajcc_pathologic_n_N1b | Lymph node pathology |
| ajcc_pathologic_n_N1c | Lymph node pathology |
| ajcc_pathologic_n_N1mi | Lymph node pathology |
| ajcc_pathologic_n_N2 | Lymph node pathology |
| ajcc_pathologic_n_N2a | Lymph node pathology |
| ajcc_pathologic_n_N3 | Lymph node pathology |
| ajcc_pathologic_n_N3a | Lymph node pathology |
| ajcc_pathologic_n_N3b | Lymph node pathology |
| ajcc_pathologic_n_N3c | Lymph node pathology |
| ajcc_pathologic_n_NX | Lymph node pathology |

Table 9: ConSurv-Rule Concept 5, Rank 5 for Run 6

| **Features** | **Alternative Name** |
|---|---|
| ENSG00000186847.6 | KRT14 |
| ENSG00000248144.6 | ADH1C |
| ENSG00000263426.2 | RN7SL471P |
| ENSG00000139329.5 | LUM |
| ENSG00000211890.4 | IGHA2 |
| ENSG00000197249.14 | SERPINA1 |
| ENSG00000143248.13 | RGS5 |
| ENSG00000198899.2 | MT-ATP6 |
| ENSG00000171428.15 | NAT1 |
| ENSG00000170345.10 | FOS |
| ENSG00000096088.16 | PGC |
| ENSG00000171346.16 | KRT15 |
| ENSG00000136881.12 | BAAT |
| ENSG00000136929.13 | HEMGN |
| ENSG00000196296.14 | ATP2A1 |
| ENSG00000211662.2 | IGLV3-21 |
| ENSG00000164404.8 | GDF9 |
| ENSG00000129988.6 | LBP |
| ENSG00000167757.14 | KLK11 |
| ENSG00000134827.8 | TCN1 |
| ENSG00000253755.1 | IGHGP |
| ENSG00000141750.7 | STAC2 |
| ENSG00000135413.9 | LACRT |
| ENSG00000243302.3 | AC018638.4 |
| ENSG00000109072.14 | VTN |
| ENSG00000086967.10 | MYBPC2 |
| ENSG00000170421.12 | KRT8 |
| ENSG00000147256.12 | ARHGAP36 |
| ENSG00000131771.14 | PPP1R1B |
| ENSG00000124102.5 | PI3 |
| ENSG00000145192.13 | AHSG |
| ENSG00000265972.6 | TXNIP |
| ENSG00000163220.11 | S100A9 |
| ENSG00000133048.13 | CHI3L1 |
| ENSG00000159388.6 | BTG2 |
| ENSG00000231500.7 | RPS18 |
| ENSG00000263639.7 | MSMB |
| ENSG00000198786.2 | MT-ND5 |
| ENSG00000265929.1 | MIR5195 |
| ENSG00000211976.2 | IGHV3-73 |
| days_to_birth | Age |

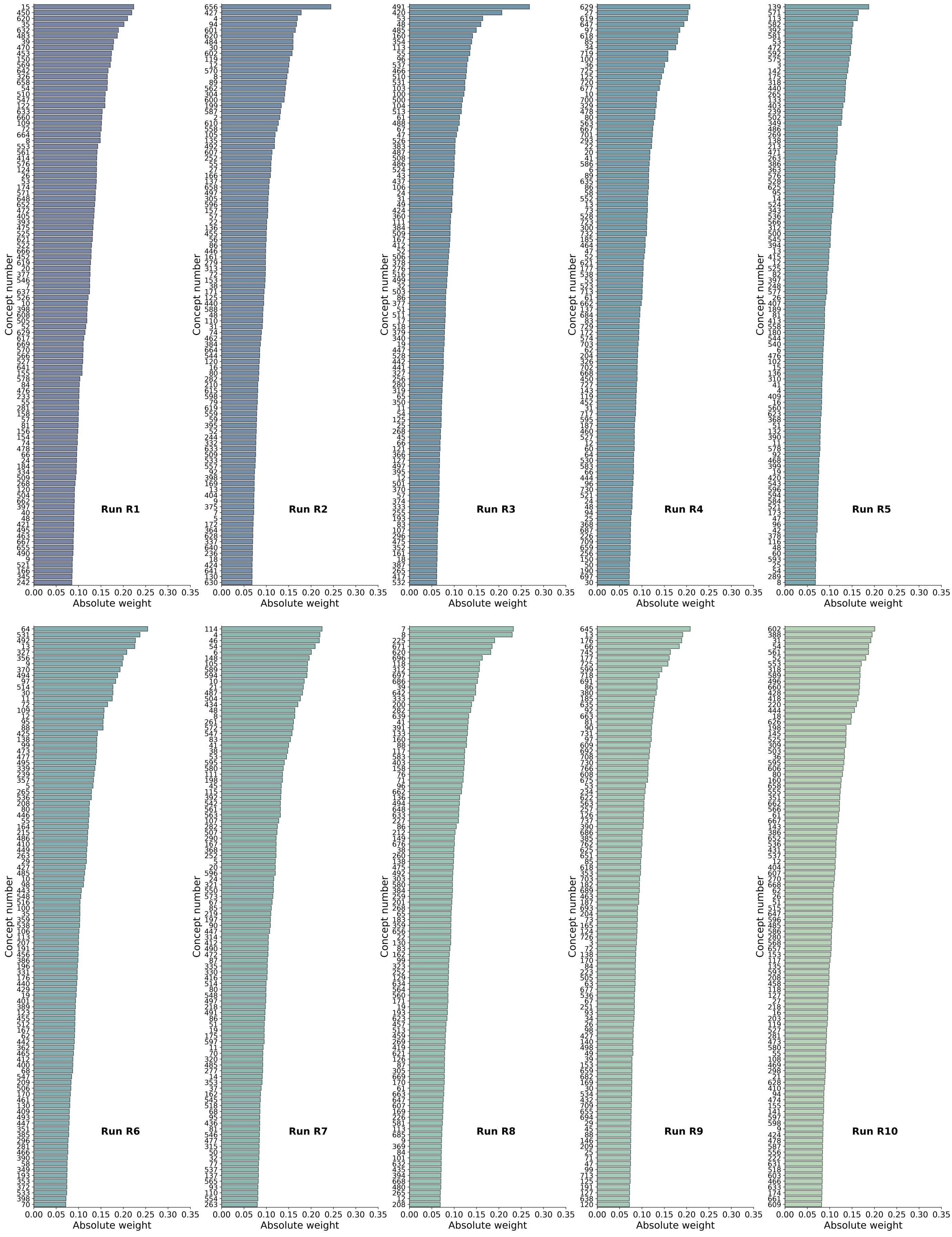


Figure 2: Top 100 concepts ordered by absolute weights in ConSurv-XGB-all across all 10 runs

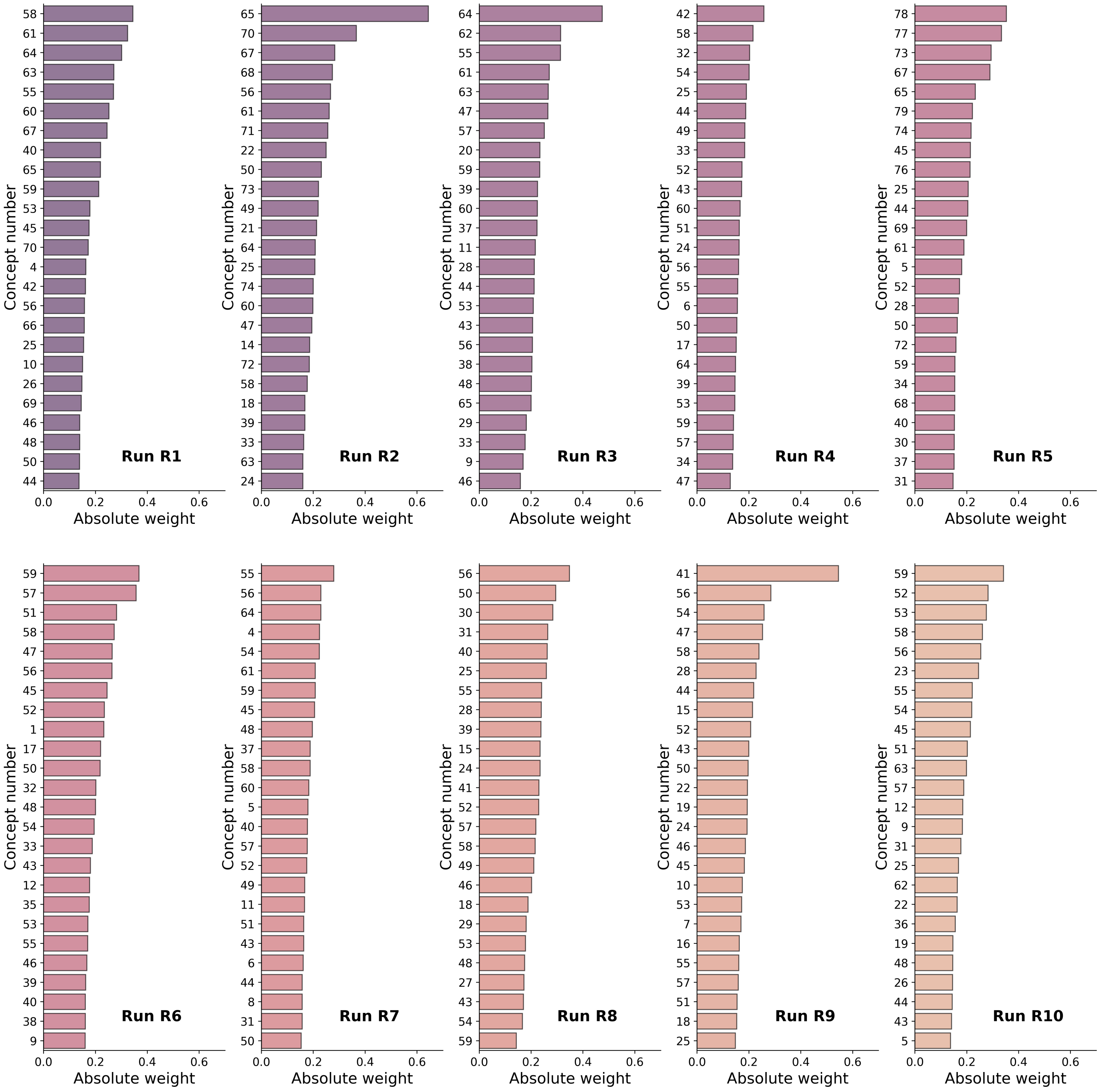


Figure 3: Top 25 concepts ordered by absolute weights in ConSurv-Rule-all across all 10 runs

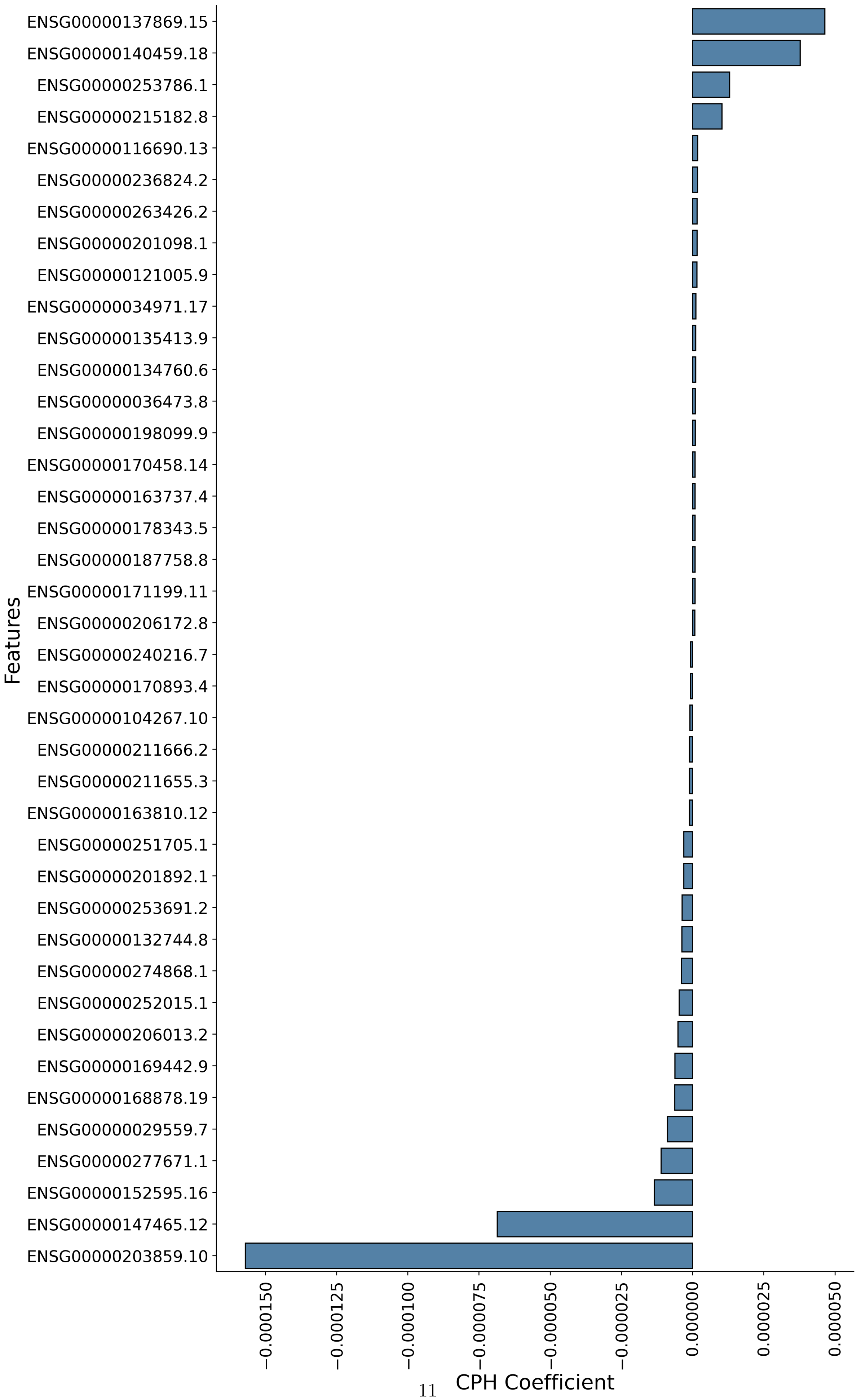




Figure 4: CPH feature coefficients for top 40 features for Run 6

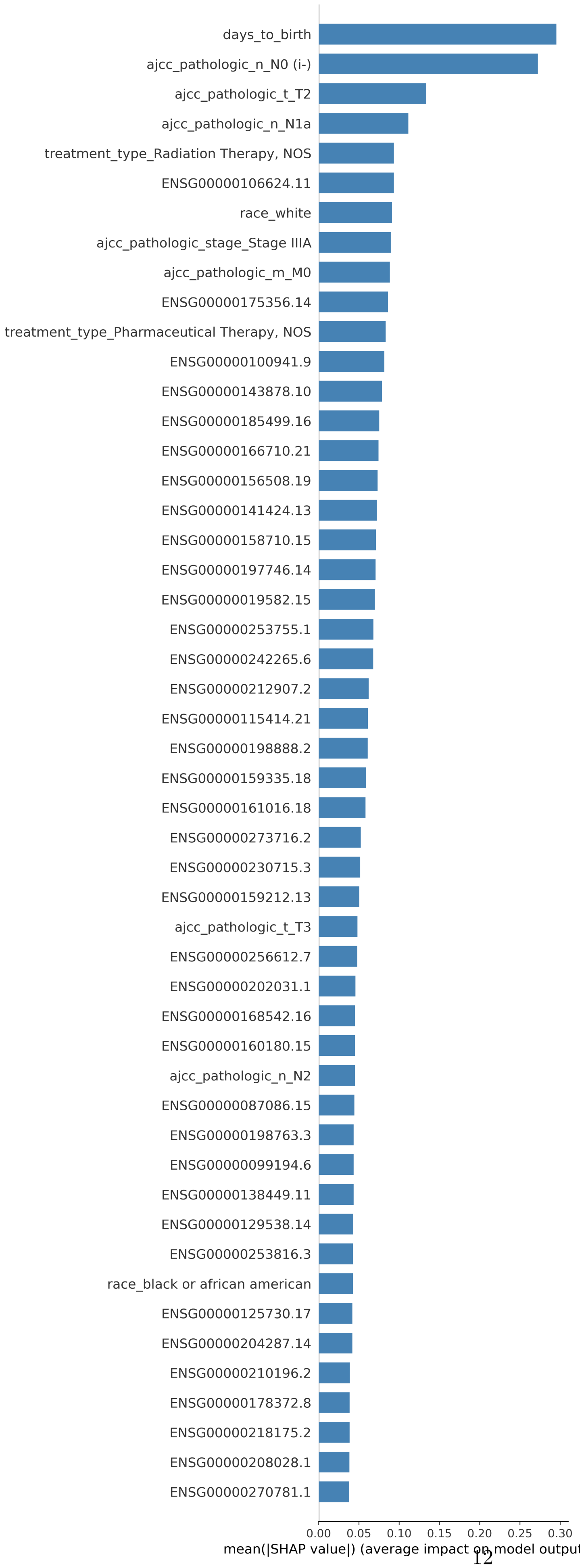




Figure 5: SHAP summary plot for DeepSurv for Run 6

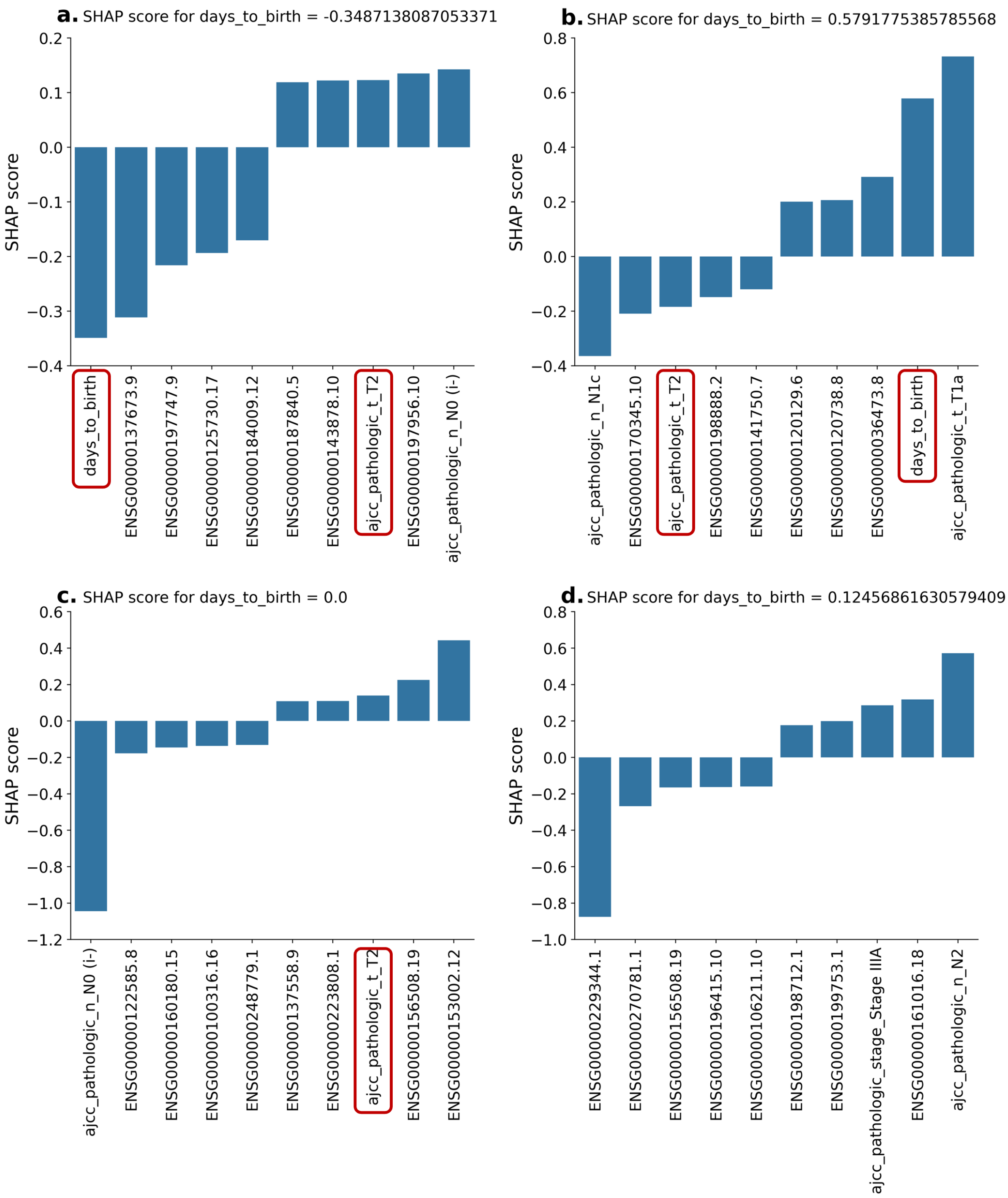


Figure 6: SHAP scores for top 10 features for four test patients for Run 6 using DeepSurv model
